# Pore structure controls stability and molecular flux in engineered protein cages

**DOI:** 10.1101/2021.01.27.428512

**Authors:** Lachlan S. R. Adamson, Nuren Tasneem, Michael P. Andreas, William Close, Eric N. Jenner, Taylor N. Szyszka, Reginald Young, Li Chen Cheah, Alexander Norman, Hugo I. MacDermott-Opeskin, Megan L. O’Mara, Frank Sainsbury, Tobias W. Giessen, Yu Heng Lau

## Abstract

Protein cages are a common architectural motif used by living organisms to compartmentalize and control biochemical reactions. While engineered protein cages have recently been featured in the construction of nanoreactors and synthetic organelles, relatively little is known about the underlying molecular parameters that govern cage stability and molecular flux through their pores. In this work, we systematically designed a 24-member library of protein cage variants based on the *T. maritima* encapsulin, each featuring pores of different size and charge. Twelve encapsulin pore variants were successfully assembled and purified, including eight designs with exceptional and prolonged thermal stability. While pores lined with negatively charged residues resulted in more robust assemblies than their corresponding positively charged variants, we were able to form stable assemblies covering a full range of pore sizes and charges, as observed in seven new cryo-EM structures of pore variants elucidated at resolutions between 2.5-3.6 Å. Alongside these structures, molecular dynamics simulations and stopped flow kinetics experiments reveal the importance of considering both pore size and surface charge, together with flexibility and rate determining steps, when designing protein cages for controlling molecular flux.

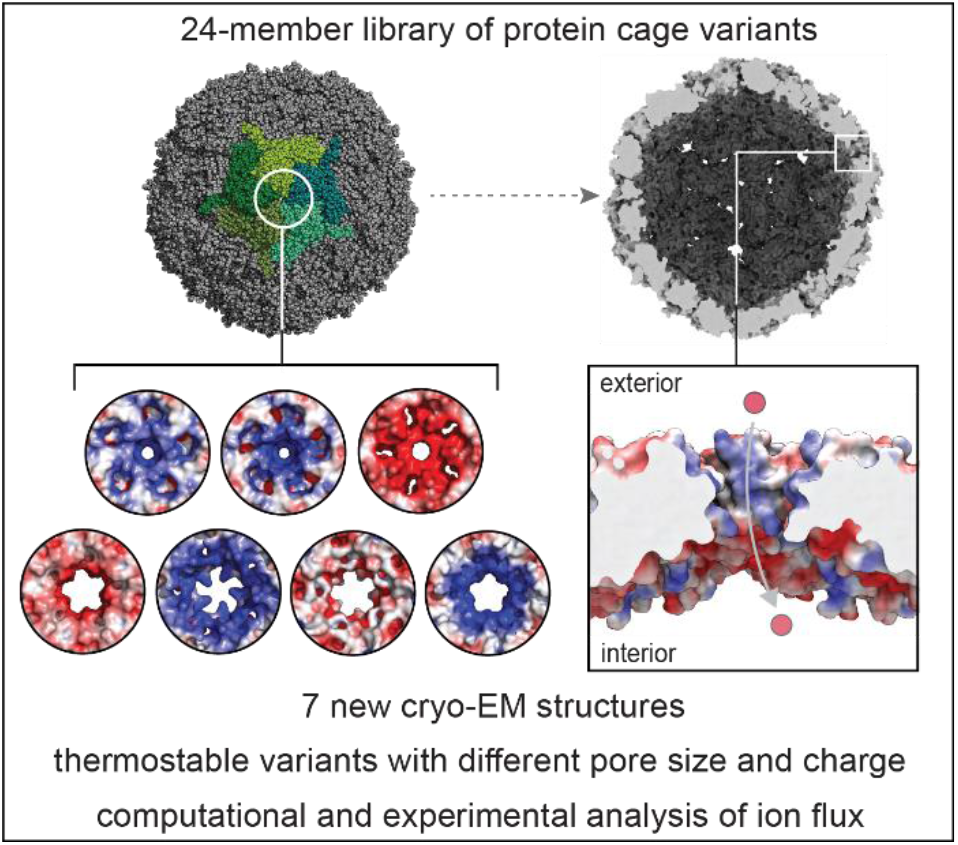

## Introduction

Spatial organization is a fundamental principle used by Nature to coordinate the complex network of biochemical reactions necessary for life. In recent years, protein cages have emerged as a diverse and widespread architecture for the organization of biochemical processes in bacteria, alongside canonical lipid-bound organelles and phase-separated membraneless condensates.^1^ An illustrative example of protein cage function is the carboxysome, a nanoscale compartment that facilitates carbon fixation in cyanobacteria and other autotrophs.^2^ The proteinaceous outer shell of a carboxysome is thought to function as a semi-permeable barrier, permitting influx of the substrate HCO_3_^-^, while retaining the volatile intermediate CO_2_ and excluding the entry of cross-reactive O_2_.^3–7^ This control over molecular flux afforded by the outer cage is crucial for achieving optimal reaction kinetics.

Inspired by the carboxysome and other naturally-occurring protein cages, chemists and bioengineers have endeavored to use modified and *de novo* protein cages as nanoreactors, hosting custom chemical reactions within their controlled internal environment.^8–14^ Despite many efforts, the design of customizable nanoreactors remains in its infancy, as the impacts of restricted molecular flux into and out of encapsulated systems are still poorly understood. Numerous groups have reported the successful encapsulation of enzymes within protein cages,^12,15–20^ but these engineered cages have yet to emulate or surpass natural systems in terms of controlling substrate influx, co-factor retention, and product efflux. Seminal work by Tullman-Ercek and co-workers indicated the possibility of charge-based control, using anionic point mutations at the pores of MS2 bacteriophage capsid particles to slow the efflux of negatively charged species, based on modelling of experimentally-determined enzyme activity measurements.^21^ Meanwhile, Lutz and co-workers have instead reported pore size as a crucial determinant of molecular flux into encapsulin cages.^22^ Recently, Douglas and co-workers used dendrimer-conjugated NADH to study molecular flux into P22 virus-like particles, studying trends in encapsulated enzyme activity upon varying the size and charge properties of the dendrimer.^23^

While these and other studies^24–27^ suggest the tantalizing possibility of programmable molecular flux through a combination of size and charge, progress in nanoreactor design continues to be limited by the lack of systematic *in vitro* studies that directly quantify flux events through cage pore variants. The independent contributions of size and charge cannot be easily deconvoluted through indirect experiments on a limited set of cages. Furthermore, insights into the impact of pore design have been limited by the lack of elucidated structures for pore-engineered cages in earlier studies.

In this work, we investigate the structure, stability, and porosity of engineered cages from a 24-member library covering different pore sizes and surface charge profiles, each based on the encapsulin protein from *T. maritima*.^28^ Encapsulins are a family of prokaryotic protein organelles that self-assemble into virus-like capsid structures of icosahedral symmetry, with external diameters ranging from 24-42 nm depending on the species.^28–33^ Our rationale for choosing encapsulins is their ability to form extremely robust and homogeneous cage assemblies from a single type of protein building block. The *T. maritima* encapsulin features narrow 3 Å pores at the five-fold axes of symmetry lined by a flexible peptide loop that Lutz and co-workers have shown is amenable to mutation and deletion (Figure 1a).^22^ The combination of robustness and engineerability has given rise to a growing number of literature reports where encapsulins have been used in biotechnological applications such as synthetic organelles, drug delivery, and vaccines.^12,34–39^

**Figure 1.**
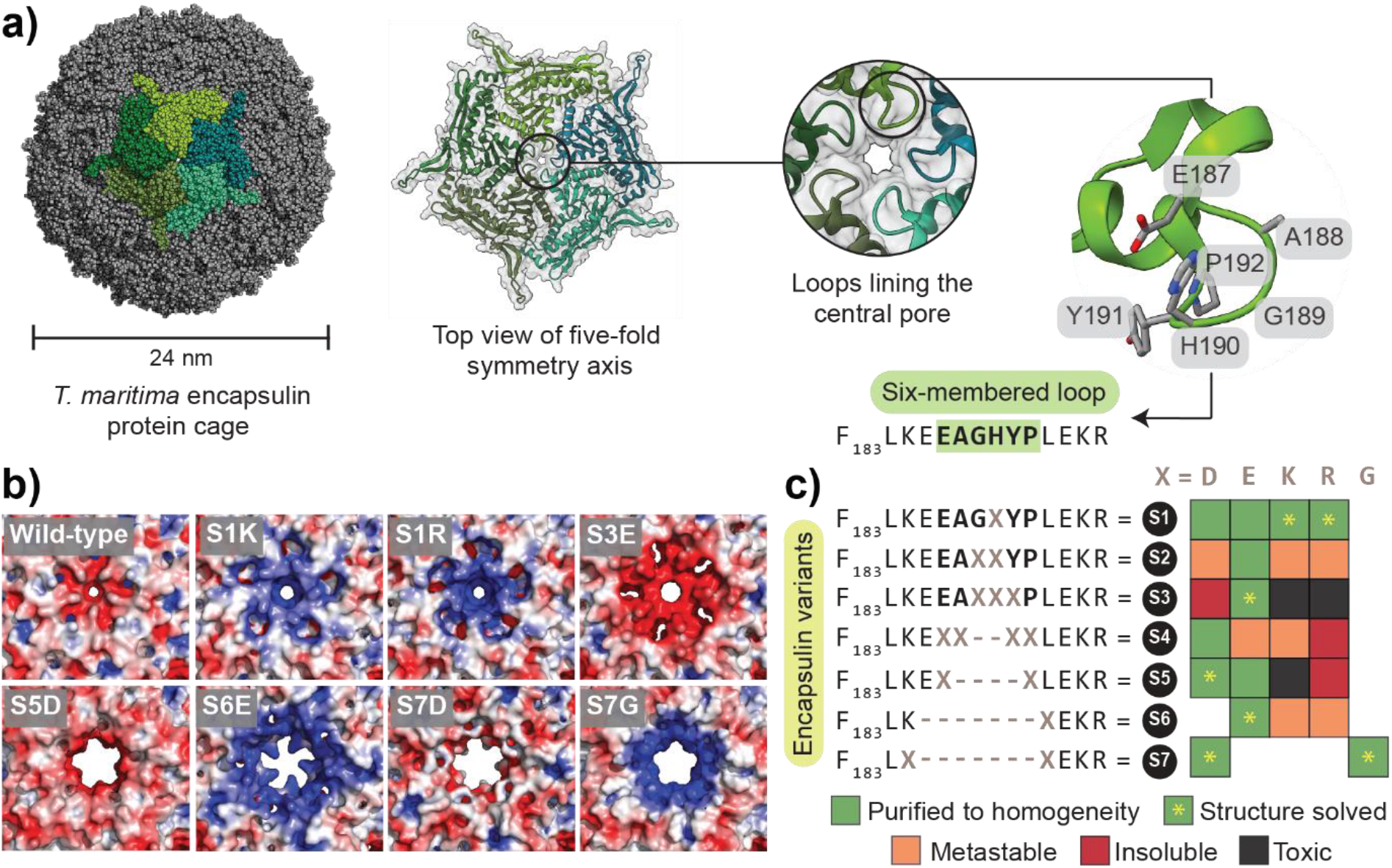
Sequence, structure and solubility of the encapsulin pore mutant library. **a)** The self-assembled hollow cage structure of the *T. maritima* encapsulin with an external diameter of 24 nm (structure from PDB 3DKT).^28^ Top view shows the location of central pore at five-fold symmetry axis of *T. maritima* encapsulin, lined by peptide loops from each monomer (residues 187-192). **b)** Novel structures of seven encapsulin variants that we determined by cryo-EM, featuring diverse pore surface electrostatics that differ from the wild-type cage. **c)** Matrix displaying 24 encapsulin variants **S1X**-**S7X** we designed in this study, along with the **S7G** positive control,^22^ indicating their sequence design and solubility post-expression and during purification. Metastable indicates precipitation during purification, while toxic indicates impaired growth of the expression host.

Herein, we systematically determine the impact of pore structure and charge on nanoreactor design. We test the integrity of a homogeneously purified encapsulin pore variant library using a combination of negative stain transmission electron microscopy (TEM), size-exclusion chromatography coupled to multi-angle light scattering (SEC-MALS), and thermal denaturation coupled to dynamic light scattering (DLS). We then characterize the size and surface electrostatics of our pore designs by determining structures of seven pore variants by single particle cryo-electron microscopy (cryo-EM), and quantify the impact of size and charge on molecular flux using a combination of molecular dynamics simulations and stopped-flow luminescence spectroscopy in an established lanthanide binding tag (LBT) assay^22,40^.

## Results

### Design and purification of charged pore variants

We designed 24 novel variants of the *T. maritima* encapsulin (**S1X**-**S7X** where X=D/E/K/R, Figure 1b) by introducing a combination of charged point mutations and deletions at the flexible loop lining the five-fold symmetrical pore. This pore region had previously been reported by *Williams et al*. to be partially tolerant to mutagenesis with alanine or glycine without completely compromising cage self-assembly.^22^ To systematically explore the effects of positively and negatively charged substituents in tandem with pore-widening deletions, we incorporated aspartic acid, glutamic acid, lysine, and arginine as single, double, and triple point mutants (**S1**-**S3**), and as flanking positions adjacent to deletions of two, four, or seven residues (**S4**-**S7**). As a positive control, we also included a pore deletion flanked with glycine (**S7G**) that had previously been reported to increase molecular flux.^22^

All encapsulin variants were recombinantly expressed in *E. coli* and subjected to a multi-step purification protocol to obtain pure and monodisperse encapsulin assemblies. This protocol involved initial enrichment from cell lysate by precipitation with poly(ethylene)glycol followed by crude size-exclusion chromatography (SEC), ion-exchange chromatography (IEC) to remove surface-bound nucleic acids, and final polishing with two further rounds of SEC (SI Section 3).

### Charge of pore residues governs robustness and in-solution stability of cage assembly

We found that eleven new encapsulin pore variants and the previously reported positive control **S7G** could be expressed and purified to homogeneity (Figure 1c). An additional seven variants were metastable, eluting at the expected size in the first round of SEC, but precipitating irreversibly after fractionation or during subsequent purification steps. Of the remaining six variants, **S3D, S4R** and **S5R** were expressed in an insoluble form (SI Figure S3.1), while **S3K, S3R** and **S5K** resulted in toxicity for the expression host.

Negatively charged variants were clearly more capable of forming assembled cages than their corresponding positively charged variants. The only completely insoluble negatively charged variant was the aspartic acid triple point mutant **S3D**, possibly due to a combination of unfavorable charge repulsion and steric effects. Intriguingly, the analogous glutamic acid mutant **S3E** formed soluble assemblies, potentially due to the extra methylene unit allowing greater flexibility to relieve geometric and charge-repulsion constraints. In stark contrast, the only two positively charged variants that we could purify to homogeneity were the single point mutants **S1K** and **S1R**.

### Assembled pore variants retain native structure and high thermal stability

All successfully purified assemblies retained a similar overall molecular architecture to that of the wild-type *T. maritima* encapsulin (Figure 2a; SI Section 4). Both TEM and DLS data showed that our purified samples were monodisperse, confirming the homogeneity of cage assembly. The expected diameter of 24 nm was observed for all assembled cages by TEM of negatively stained samples, with no evidence of aberrant assemblies (Figure 2ai; SI Figure S4.1). The hydrodynamic size of the cages in Tris buffer was measured by DLS and found to be consistent with the TEM data (Figure 2aii; SI Table S4.3). Purity and monodispersity were further confirmed by SEC-MALS, with a single peak observed for each purified variant (Figure 2aiii; SI Figure S4.2) giving an assembled mass of 2-2.5 MDa. SDS-PAGE analysis showed a clear band for the encapsulin monomer at the expected size of 33 kDa (Figure 2aiv; SI Figure S3.4).

**Figure 2.**
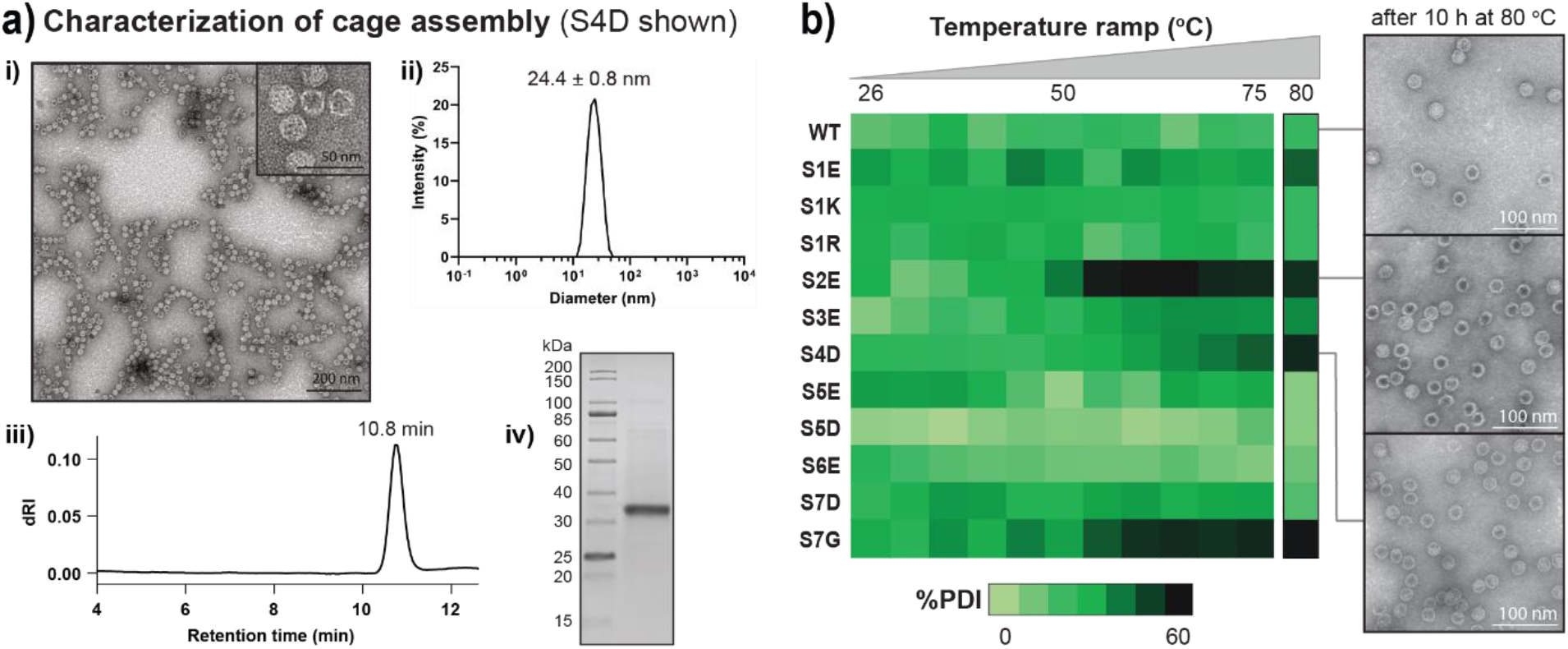
Pore variants can self-assemble and are stable at high temperatures. **a)** Exemplar characterization data for purified variant **S4D** (data for other variants can be found in SI Sections 3 and 4). (i) Negative stain TEM image of assembled cages. (ii) DLS profile showing the expected hydrodynamic size. (iii) Analytical SEC chromatogram (Bio SEC-5 1000 Å column) showing the presence of a single monodisperse species. (iv) SDS-PAGE analysis showing a single band corresponding to the encapsulin monomer. **b)** Heat map of the change in polydispersity index (%PDI) upon increase in temperature (from left to right: 26, 30, 36, 40, 44, 50, 56, 61, 66, 70 and 75 °C). TEM images of selected variants show the presence of intact assemblies after 10 h incubation at 80 °C.

Prolonged thermal stability was observed for purified samples of all but two of the new charged pore variants **S2E** and **S4D** and the positive control **S7G**, revealing a remarkable degree of structural integrity and amenability to mutagenesis (Figure 2b). Upon heating samples to 75 °C while monitoring hydrodynamic size and polydispersity by DLS, we only observed an irreversible increase in polydispersity for **S2E, S7G**, and to a lesser extent **S4D**, indicative of aggregation or denaturation. A further overnight incubation of all samples at 80 °C did not impact the stability of the assemblies. Despite the thermal instability of **S2E** and **S4D**, we still observed the presence of intact assemblies by negative stain TEM after prolonged heating, indicating that only partial denaturation or aggregation had occurred.

### Cryo-EM structures reveal that a range of pore sizes and charges can be obtained

We elucidated the atomic structures of seven selected pore variants by single particle cryo-EM (SI Figures S6.1 and S6.2), displaying a diverse range of pore sizes and surface charge profiles (Figures 3 and 4). Notably, we were able to obtain both positively and negatively charged pores in our variant library, despite the clear trend against positively charged mutants at the primary sequence level. The structures for encapsulin cages containing point mutations (**S1R**/**S1K**/**S3E**), a partial loop deletion (**S5D**), and complete loop deletion (**S6E**/**S7D**/**S7G**), were determined at resolutions ranging from 2.5-3.6 Å. Further, all variants contained clear density for a tightly bound flavin-based molecule located close to the three-fold symmetry axis which was recently reported,^41^ but had not previously been identified in the original structural study^28^ or observed in any other T1 encapsulin (SI Figure S4.4). Pore channel dimensions were calculated based on the cryo-EM structures using the program HOLE,^42^ while electrostatic surface potentials were calculated using the software APBS using default parameters.^43^

**Figure 3.**
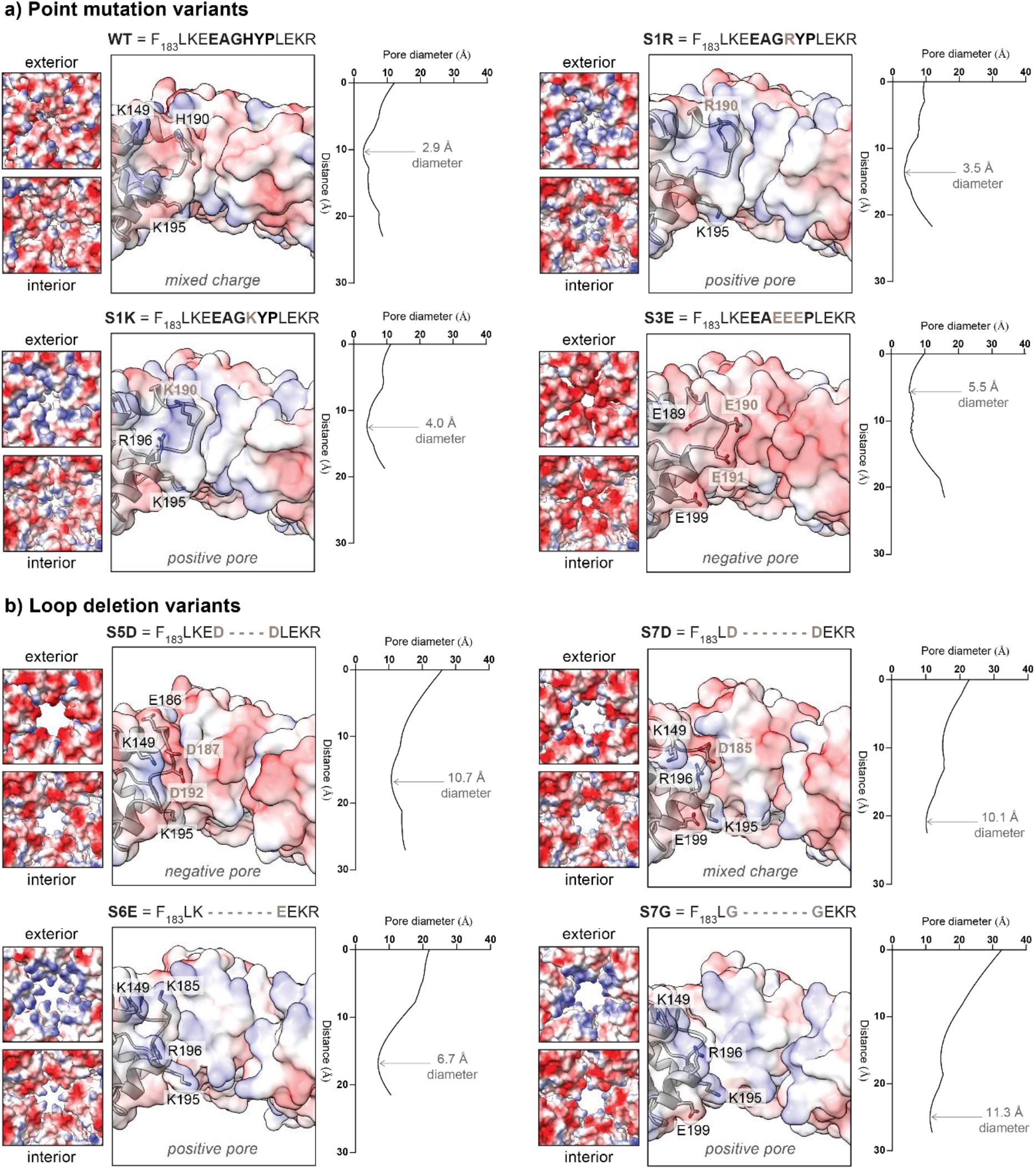
Cryo-EM structures of the pore channels show altered surface electrostatics and pore diameters. **a)** Point mutation variants (**S1R**/**S1K**/**S3E**). **b)** Loop deletion variants (**S5D**/**S6E**/**S7D**/**S7G**). For each pore channel, the exterior and interior surface charge is shown, where red is negative and blue is positive. A cross-sectional view of each pore is also shown, presenting three of the five monomers at the five-fold symmetric pore. Each corresponding graph displays the static pore size throughout the channel measured, indicating the narrowest point for each variant. Distance refers to the position along the pore axis from exterior to interior, while pore size is calculated using the program HOLE.^42^ The wild-type structure is shown for comparison (PDB 3DKT).^28^ For loop deletion variants, residue numbering is kept consistent with wild-type to facilitate comparison.

**Figure 4.**
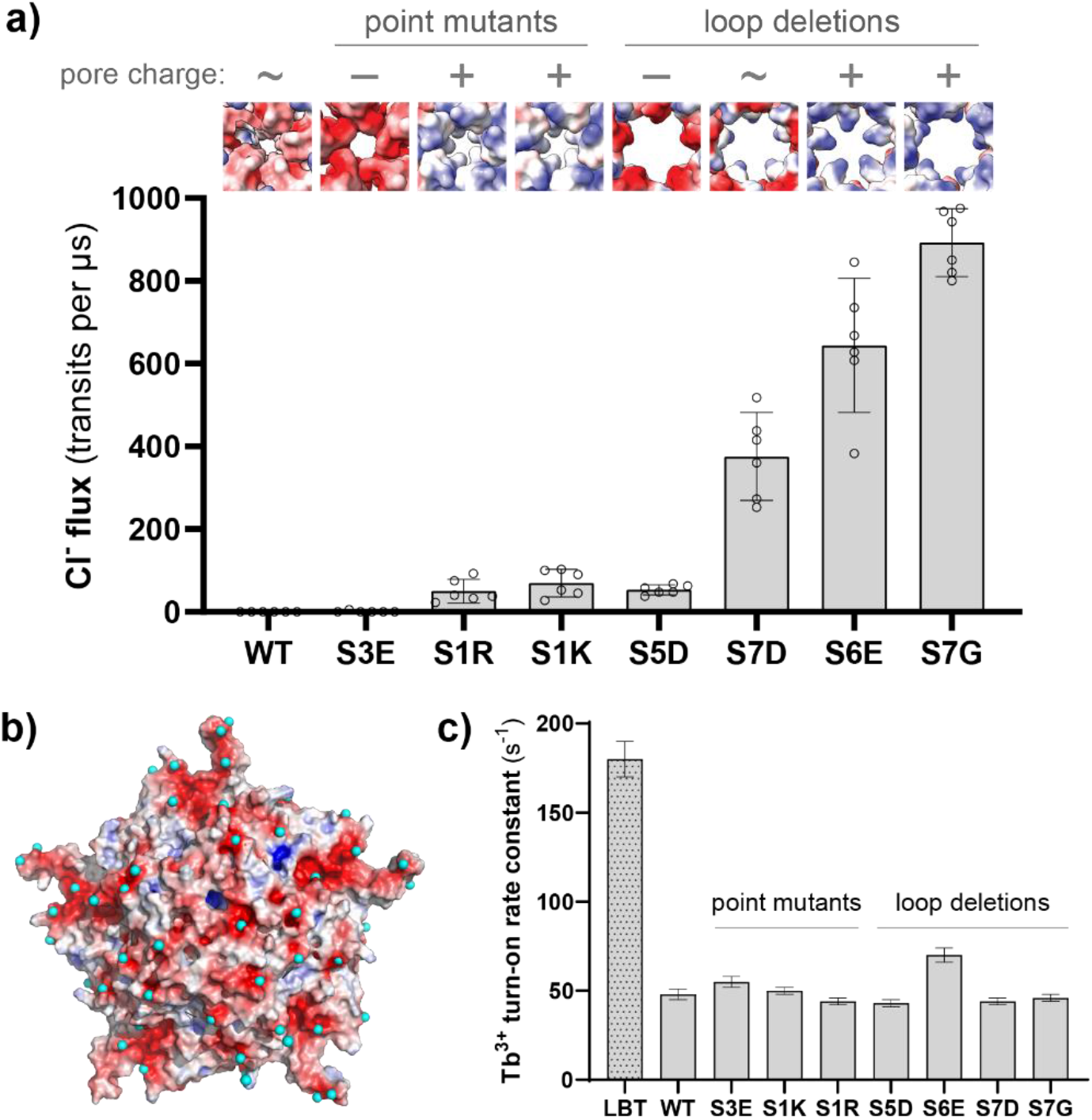
Rates of molecular flux are determined by size and charge, but surface interactions can impact flux simulations, while pore flux is not always the experimental rate determining step. **a)** The flux of chloride anions through the five-fold pore is controlled by a combination of pore size and charge, as determined by molecular dynamics simulations using the elucidated cryo-EM structures. All pairwise differences between the loop deletion variants are statistically significant in a one-way ANOVA with Tukey’s HSD test (p<0.0001). Each hollow circle represents the rate of chloride transit events through the pore during a single simulation replicate, while the bar graphs represent the mean value, and the error bars represent a 95% confidence interval. The cryo-EM exterior pore structures are shown above each bar, along with the overall charge as positive (+), mixed (∼), or negative (–). **b)** Examples of metastable Tb^3+^ binding sites on the surfaces of the encapsulin pentamer, where the ions did not move away from the surface site during the entire 400 ns simulation timeframe. One frame of the wild-type encapsulin simulation is displayed, with bound Tb^3+^ ions shown as aqua spheres, and the wild-type pentamer exterior surface shown as the charge surface, where red is negative and blue is positive. **c)** Overall rate constants for Tb^3+^ turn-on luminescence from stopped-flow kinetics experiments, alongside the rate constant for the free non-caged lanthanide-binding tag peptide (LBT), suggesting that flux through the pore is not the rate-determining step in this specific experimental assay.

The single point mutants **S1R** and **S1K** retain similar pore dimensions in comparison to the wild-type, but a complete inversion of surface electrostatics is observed at the exterior entrance of the pore (Figure 3a). The inversion to a positively charged entrance arises from the introduction of basic side-chain functional groups, together with subtle backbone conformational changes. In the wild-type encapsulin, the backbone carbonyl oxygen of H190 points directly into the pore, conrtibuting a negatively charged ring at the entrance of the pore. After mutation to R190 and K190, the peptide loop shifts outwards, such that the backbone carbonyl oxygen moves to point obliquely away from the pore lumen. No significant changes are observed on the positively charged interior surface lined by the side-chain of K195.

Major conformational shifts in the **S3E** variant results in strong negative charge throughout the pore channel and both entrances (Figure 3a). The negative charge arises from E190 and E191, which line the external and internal entrances of the pore respectively as expected, while the backbone carbonyl oxygen of E190 also points directly into the pore. The large 90° sideways shift in the pore loop conformation enables the increased steric bulk of the glutamate sidechains to be accommodated, resulting in an increase in overall pore diameter to 5.5 Å.

The deletion variant **S5D** removes most of the pore loop, widening the pore to a diameter of 10.7 Å (Figure 3b). The loop between the two helices leading to and away from the pore is replaced by a short pseudo-helical turn spanning residues 186/187/192/193 (residues 188-191 are deleted), with side-chain carboxylates of D187 and D188 pointing directly into the pore. The net result is a strongly negative pore channel and exterior entrance, while K195 continues to contribute some positive charge at the interior entrance.

The removal of three more residues in the deletion variants **S6E**/**S7D**/**S7G** converts the loop into a short linker, further widening the pore exterior entrance (Figure 3b). Unexpectedly, **S7G** and **S6E** both have positive pores despite the incorporation of neutral and negative residues respectively. The non-charged double glycine mutation of the **S7G** variant designed by *Williams et al*.^22^ allows the pore-lining residues K195 and R196 to dominate the electrostatic profile, leading to a highly positive and concave pore opening and channel from the exterior viewpoint. On the interior surface, E199 contributes some negative charge around the interior opening, but this is balanced by the positive charge of K195. Comparing variants **S7G** and **S6E**, introduction of K185 and E193 in place of two glycines leads to a similar electrostatic profile, where K185 provides positive charge in addition to K149, while E193 is buried and does not contribute significantly to the charge within the pore channel. In contrast, the introduction of D185 in variant **S7D** inverts the electrostatics of the exterior entrance while partially neutralizing the pore channel itself.

We note that in all deletion variants, the flexible side-chain of K195 is shown in a conformation that points into the pore in our structural models. As the HOLE program relies on the static structures provided, the calculated pore width at the interior entrance is not expected to be precise, especially for **S6E** where we anticipate a wider pore diameter after accounting for K195 side-chain flexibility. Indeed, conducting HOLE calculations during molecular dynamics simulations on the **S6E** pore confirms that this pore is similarly sized to the other loop deletion mutants **S7D** and **S7G** (SI Figure S8.2).

To investigate pore residue flexibility, we generated local resolution estimates for our final cryo-EM maps using cryoSPARC and an FSC gold-standard threshold of 0.143. It is well established that a local decrease in resolution often correlates with increased local flexibility of loops and side chains.^44^ Based on our analysis (SI Figure S6.3), all pore mutants display locally decreased resolutions between 0.3 and 0.5 Å in the residues surrounding the pore exterior. All loop deletion variants (**S5D**/**S6E**/**S7D**/**S7G**) and the triple point mutation variant **S3E** show a pronounced local decrease in resolution for all residues lining the interior of their wide pores including K195. On the other hand, the local resolution of the pore interior for the single point mutation variants **S1K** and **S1R** seems to represent a local resolution maximum. For **S1K** and **S1R**, residue Y191 represents the narrowest point of the pore and shows a local increase in resolution of ca. 0.5 Å indicating a locally rigid pore. Our local resolution analysis suggests that loop deletion and variants with increased negative charge density lead to generally more flexible pore residues. These trends in pore flexibility were confirmed by molecular dynamics simulations, where the broader distribution of pore sizes was observed for the loop deletion variants (SI Figure S8.2).

### Molecular dynamics reveal the effects of pore size and charge on molecular flux

With cryo-EM structures in hand, we used molecular dynamics simulations to quantify the impact of pore variation on molecular flux, finding that both size and charge complementarity play a critical role in controlling flux. As a proxy for quantifying molecular flux, we tracked the diffusion of chloride ions through the pore variants during simulations of encapsulin pentamers. The total number of chloride flux events through the five-fold pore of each variant was tallied over six replicate 400 ns simulations, revealing significant differences between each variant.

Although widening the pore size generally resulted in higher rates of molecular flux as expected, pore charge had an equally important role in controlling flux (Figure 4a, Supplementary Videos 1 and 2). Across the similarly-sized loop deletion variants **S5D**/**S6E**/**S7D**/**S7G**, there was a clear dependence on charge complementarity, with positively charged **S6E** and **S7G** pore displaying the greatest flux of chloride anions, the partially neutralized **S7D** pore showing moderate flux, and the highly negative **S5D** pore showing low flux similar to that of the narrower point mutants **S1R** and **S1K**. This result demonstrates that charge repulsion can override the effect of pore widening. Similar trends were observed across the point mutants, with the positive **S1R** and **S1K** pores having significantly higher rates of chloride flux than the negative **S3E** pore and the wild-type pore with a mix of partial positive and negative charge.

In contrast to anion flux, our simulations did not provide sufficient data to quantify cation flux on pore variants. We chose terbium(III) as the chloride counterion, matching the TbCl_3_ used in the lanthanide-based luminescence assay previously reported by Williams *et al*. for measuring the kinetics of ion influx into encapsulins.^22^ On the simulation timescale, no Tb^3+^ flux events were observed in most replicates for all pore variants, as a significant proportion of Tb^3+^ ions were bound throughout the entire simulation at negative patches on both the interior and exterior encapsulin surfaces (Figure 4b, Supplementary Video 3). Interestingly, this behavior is reminiscent of Fe^2+^ binding to surface-exposed carboxylates during flux within ferritin cages.^45^ While the impact of surface binding is expected to vary with ionic strength and relative stoichiometry, there is a high ratio of encapsulin to Tb^3+^ in the simulations, necessitated by the dimensions of the simulation system, resulting in a lack of free Tb^3+^ available for pore flux.

### Overall reaction kinetics depends on whether influx or cargo kinetics is rate determining

The overall kinetics of compartmentalized systems are complex, with a dependence on the relative rates of molecular flux through the cage pores as well as the inherent kinetics of the encapsulated cargo.^46^ Upon conducting terbium luminescence experiments using stopped-flow kinetics in accordance with Williams *et al*. (see SI Section 7 for details) on our fully-characterized subset of pore variants,^22^ we observed unexpected experimental trends which strongly suggest that luminescent complex formation is in fact the rate-determining step in this specific caged system, rather than molecular flux through pores.

When measuring the turn-on kinetics of excess Tb^3+^ interacting with a lanthanide-binding tag (LBT) in the form of a free peptide in solution, maximal luminescence signal corresponding to equilibrated complexation of Tb^3+^ was reached in ∼30 ms, giving a pseudo-first order rate constant of 180±10 s^-1^ (Figure 4c). In comparison, the encapsulated version of the LBT *N*-terminally fused inside wild-type encapsulin showed four-fold slower luminescence turn-on kinetics with a pseudo-first order rate constant of 48±3 s^-1^, thus matching the trend observed in the literature.^22^ However, this modest drop in rate constant is not necessarily indicative of the encapsulin shell as a rate-determining barrier to diffusion, as for example, competition between multiple encapsulated LBTs within the encapsulin lumen could also result in complex kinetics that can lower the apparent first-order rate.^47,48^

Furthermore, all pore variants exhibited relatively similar slow rate constants for Tb^3+^ turn-on kinetics (Figure 4c, SI Figure S7.1), suggesting that the overall kinetics may primarily be governed by Tb^3+^-LBT luminescent complex formation as a rate-determining step, rather than transit through the pores. Apart from **S3E** and **S6E** which showed slight increases over the wild-type rate with pseudo-first order rate constants of 55±3 s^-1^ and 70±4 s^-1^ respectively, there were minimal differences in rate for the remainder of the variants, including the **S7G** control featuring a wide but positively-charged five-fold pore (46±2 s^-1^), which had previously been reported to give a seven-fold increase in Tb^3+^ flux using a comparable experimental setup.^22^

Upon estimating the theoretical rates of Tb^3+^ diffusion through the wild-type encapsulin based on two models, one based on the permeability of encapsulin pores^46,49^ and another on Tb^3+^ depletion around the encapsulin shell (SI Section 9),^50^ the estimated theoretical rate constant in both scenarios (∼10^4^ s^-1^ and ∼10^6^ s^-1^) was several orders of magnitudes greater than the measured rate constant for Tb^3+^-LBT complex formation (200 s^-1^). While the theoretical diffusion rates do not account for any molecular interactions between Tb^3+^ and the encapsulin pore which may decrease the rate constant, the magnitude of the discrepancy between estimated diffusion rates and the experimentally measured rate constant of luminescent complex formation suggests that differences in diffusion rates cannot be detected using this experimental setup.

While Tb^3+^ was chosen for its ability to be quantified directly *via* luminescence in stopped-flow experiments, a more realistic nanoreactor scenario would involve the flux of small molecules such as reaction substrates. In these more realistic cases, we expect to see the size- and charge-dependent behaviors observed in the chloride ion simulations, due to the slower rate of flux through the pore, but only when the functional readout (eg. enzymatic turnover) is faster than this diffusional flux. Indeed, recent enzymatic measurements from the same research group using the **S7G** control variant provide indirect evidence that small molecule flux is enhanced by widened five-fold pores.^51^

## Discussion

In our study, we show that it is possible to build a library of protein cages featuring pores of different size and charge, showcasing the remarkable robustness of encapsulins and their tolerance to mutation, even when subjected to extended periods of high temperature. Our cryo-EM structures provide atomic-level understanding of how the designed sequence modifications translate to pore size and charge differences in the final cage assembly.

The clear preference in stability and assembly robustness towards negatively charged pore mutations in our encapsulin system has significant broader implications for protein cage design. As a general concept, it may be possible in the future to use pore charge modifications to control the integrity of cage assembly, as potentially inducible destabilization of protein cages through switching of pore charge could be applied to the controlled release of encapsulated drug cargo, or switchable assembly of nanoreactors.^33^ Furthermore, destabilization of capsids featuring pore arginine residues is a native feature of the HIV capsid, where polyanionic metabolites are required to induce stabilization of the overall assembly.^52–55^ Our encapsulin variant library provides a robust and engineerable model for studying and mimicking the behavior of these complex biological systems.

Through our cryo-EM structures, we also discover that it is still possible to generate stable assemblies featuring positively charged pores, without directly introducing new lysine or arginine residues at the pore. In particular, variants **S6E** and **S7G** surprisingly both present positively charged pore exteriors, arising from changes in backbone alignment and the influence of more remote residues, highlighting that in addition to the primary sequence, structural information is critical for understanding pore design.

The systematic trends in chloride flux between the simulated five-fold pore variants demonstrates the importance of assessing the combined impact of pore charge and size as two equally important molecular parameters for controlling flux. For pores with similar charge such as **S1R** and **S7G**, the loop deletion and subsequent pore widening results in a 20-fold difference in flux. However, for similarly sized pores such as **S5D** and **S7G**, the inversion of pore charge leads to a 16-fold difference in flux. Given the prevalence of charged functionalities in biological chemistry, such as phosphates and amino acids, due consideration must be given to both size and charge when designing nanoreactors. Combined size and charge effects were recently reported in P22 bacteriophage capsid virus-like particles, although using much larger dendrimer-based molecules to match the much more porous P22 cage.^23^

The discrepancy between our findings and those of Williams *et al*.^22^ (especially control variant **S7G**) emphasizes the challenges when quantifying kinetics in encapsulated systems. As the kinetic assay conditions themselves were closely matched, differences may have arisen during sample purification. In our hands, we found that low-resolution SEC was insufficient to obtain a contamination-free sample. Non-specifically surface bound nucleic acids have been noted as a co-purified contaminant,^56^ which we were only able to remove upon IEC, as determined by UV absorbance ratios at 260 and 280 nm. The presence of such impurities may not be observable when using other characterization methods such as SDS-PAGE and TEM. Terbium stock preparation may also have resulted in discrepancies, as non-acidic aqueous solutions degrade over time by hydroxide formation.

Ultimately, while Tb^3+^ is not an optimal choice for directly measuring pore flux, the experiments highlight the critical importance of considering the relative rates of all the mechanistic steps when designing encapsulated nanoreactor systems. From a mechanistic standpoint, due to the relatively narrow protein pores with potential transient interaction sites for Tb^3+^,^57^ we do expect to observe ion transport behavior reminiscent of ion channels such as gramicidin A,^58^ rather than simple Fickian diffusion trends as previously reported.^22^ Indeed, there is evidence that iron transport behavior in ferritin cages is facilitated by the pore structure, involving charge-selective interactions and multiple coordination sites formed by the pore residues.^45,59–62^ Nevertheless, interactions between Tb^3+^ and the pores would need to slow diffusion by several orders of magnitude for the relative rates of diffusion to match those of complex formation, which is an unlikely scenario. More plausible contributing factors for the slowed but relatively uniform turn-on kinetics across pore variants include competition between co-encapsulated LBTs which may affect binding kinetics and affinity, steric constrains on the encapsulated LBT peptide from crowding or interactions with the interior surface, and potential quenching events between neighboring LBT sites.

Along with the set of structurally characterized protein cage variants from this work, our newfound knowledge of how size and charge affect flux will enable systematic structure-based design of nanoreactor porosity for a broad range of encapsulated chemical and enzymatic reactions. Future work will involve quantifying the impact of pore structure on the conversion of enzyme substrates of different sizes and charges. In this nanoreactor context, we expect that substrates, products, and co-factors will all experience a greater diffusional barrier than a simple metal ion such as Tb^3+^, slowing the rate of molecular flux to such an extent that it plays an important role in the rate equation for the overall transformation.^46^ Our work therefore represents a crucial step in the development of programmable large-scale and multiplexed nanoreactors for simultaneously conducting otherwise incompatible reactions inside live cells.

## Experimental Methods

### Molecular cloning

Codon optimized genes for the wild-type *T. maritima* encapsulin with and without an *N*-terminal lanthanide binding tag were commercially purchased as gBlocks (Integrated DNA Technologies) and cloned into pBAD-HisA vector using the BglII and HindIII restriction sites. The untagged pore variant plasmids were generated by whole-plasmid PCR with mutagenic primers (SI Table S2.1), using untagged wild-type *T. maritima* plasmid as a template followed by DpnI digestion and clean-up, and subsequent transformation of the cleaned linear PCR products directly into *E. coli* strain DH5α. The tagged pore variant plasmids were constructed by Gibson assembly using NEBuilder HiFi DNA Assembly Master Mix (NEB), where the variant encapsulin genes were individually amplified by PCR from the untagged pore variant plasmids, and the tag-containing backbone was amplified from the tagged wild-type plasmid. Chemically competent *E. coli* strain DH5α was used for all cloning procedures. Sequences of plasmids were confirmed by Sanger sequencing conducted through the Ramaciotti Centre for Genomics (UNSW, Australia). See associated GenBank files on GitHub for full sequence details (https://github.com/LauGroup/TmEnc_Flux/tree/main/plasmid%20maps).

### Protein expression and purification

Designed encapsulin variants containing *N*-terminal lanthanide binding tags (SI Table S2.2) were expressed in BL21(DE3) cells. Overnight 5 mL cultures were diluted into 400 mL of fresh LB media and grown at 37 °C until OD_600_ of 0.4 was reached. Expression was induced with 0.2% arabinose and the cells were incubated at 20 °C overnight. Cells were then pelleted by centrifugation at 3900 rcf for 15 min (Eppendorf S5810R with S-4-104 rotor) and stored at -80 °C.

Frozen cell pellets were resuspended in 24 mL of lysis buffer (20 mM Tris, 100 mM NaCl, 100 µg/mL DNase, 100 µg/mL lysozyme, pH 8) and incubated on ice for 10 min. Cells were lysed using a Sonopuls HD 4050 probe sonicator with a TS-106 probe (Bandelin) using an amplitude setting of 55% and a pulse time of 8 s on and 10 s off for a total time of 11 min. The lysate was clarified by centrifugation (Eppendorf S5430R with F-35-6-30 rotor, 7197 rcf for 7 min), further clarified by syringe filtration if needed, then to the supernatant, NaCl was added to 250 mM and PEG-8000 to 10%(w/v) final concentration. The suspension was incubated on ice for 10 min then centrifuged (Eppendorf S5430R with F-35-6-30 rotor, 7197 rcf for 7 min). The supernatant was removed, and the pellet was dissolved in a total volume of 4 mL size exclusion buffer (20 mM Tris, 100 mM NaCl, pH 8).

The redissolved protein was subject to size exclusion chromatography using a Sephacryl S-500 16/60 column at a flow rate of 1 mL/min (SI Figure S3.2). Fractions containing encapsulin were identified by SDS-PAGE, pooled, and filtered to remove nucleic acids by ion exchange chromatography using a HiPrep Q XL 16/10 column at a flow rate of 5 mL/min (SI Figure S3.3), resulting in A260/A280 ratios of less than 0.8 in all cases. Flow-through fractions containing encapsulin were concentrated by ultrafiltration (Amicon 100 kDa MWCO) and subjected to two rounds of size exclusion chromatography with a Superose 6 10/300 GL column at a flow rate of 0.3 mL/min (SI Figure S3.4a). Protein concentrations were determined by absorbance at 280 nm. Purified proteins were stored short-term in size exclusion buffer at 4 °C until further use.

### Negative stain transmission electron microscopy (TEM)

Encapsulin samples for negative stain TEM were diluted to 100 µg/mL in 20 mM Tris pH 8. Gold grids (200-mesh coated with Formvar-carbon film, EMS) were made hydrophilic by glow discharge at 50 mA for 30 s. The grid was floated on 20 µL of sample for 1 min, blotted with filter paper, and washed once with distilled water before staining with 20 µL of uranyl acetate for 1 min. TEM images were captured using a JEOL-1400 at 120 keV at the Australian Centre for Microscopy and Microanalysis.

For TEM after DLS thermal stability experiments, samples were imaged with a Hitachi HT770 at 80 keV, using the same protocol except the grids were not glow discharged.

### Size-exclusion chromatography with multi-angle light scattering (SEC-MALS)

SEC-MALS was performed using a Nexera Bio-HPLC (Shimadzu). Separations were performed using a Bio SEC-5 1000 Å 7.8 × 300 mm column (Agilent) with a Bio SEC-5 2000 Å 7.8 × 300 mm guard column (Agilent). Multi-angle light scattering was measured using a Dawn 8 Multi-Angle Light Scattering detector (Wyatt Technologies). Concentration was calculated from change in refractive index assuming a 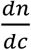 of 0.185. PBS (137 mM NaCl, 2.7 mM KCl, 10 mM Na_2_HPO_4_, 1.8 mM KH_2_PO_4_, pH 7.4) was used as the mobile phase and kept at a constant flow rate of 1 mL/min. Samples for SEC-MALS analysis were typically diluted to 200 µg/mL and 25 µL was injected with a run time of 20 min. Molecular weights were calculated using ASTRA 7.3.0 software (Wyatt Technologies).

### Dynamic light scattering (DLS)

The hydrodynamic size of encapsulin particles was determined using a Zetasizer Nano ZS (Malvern Analytical). DLS measurements were taken at 25 °C at a concentration of 17 µg/mL encapsulin in size exclusion buffer.

For thermal stability experiments, samples were loaded in duplicate on black glass-bottomed 384-well polystyrene plates (Corning) and overlayed with silicon oil (Sigma-Aldrich). Scheduled measurements were performed using a DynaPro III Plate Reader (Wyatt Technologies) and cumulants analysis was performed on 10 acquisitions per well by Dynamics V 7.9.1.4 (Wyatt Technologies).

Starting at 25 °C the temperature was increased by 2 °C each cycle (of approximately 30 minutes each) to 75 °C. Following the thermal gradient scan, a measurement was taken at 25 °C and this was followed by further overnight incubation at 80 °C, during which measurements were made every 30 minutes.

### Cryo-electron microscopy (cryo-EM) sample preparation, data collection, and data processing

Grids for cryo-EM imaging of the five encapsulin variant samples **S1K, S1R, S3E, S6E** and **S7D** were prepared by applying 3.5 μL of purified encapsulin sample diluted in 150 mM NaCl, 25 mM Tris at pH 8.0 to a concentration of 2 mg/mL to a glow-discharged (1 min at 5 mA) holey carbon grid (Quantifoil Q225-CR1.3). This was followed by plunge-freezing in liquid ethane using an FEI Vitribot Mark IV (100% humidity, blot force: 20, wait time: 0, temperature: 22 °C). Grids were clipped and stored in liquid nitrogen until data collection.

Cryo-EM movies of samples **S1K, S1R, S3E, S6E** and **S7D** were collected at the University of Michigan Life Sciences Institute on a Glacios cryo-electron microscope (Thermo Fisher Scientific) operating at 200 kV and equipped with a K2 Summit direct electron detector (Gatan). Movies were collected at a magnification of 45,000 with a calibrated pixel size of 0.98 Å using the Leginon automated data collection software,^63,64^ with an exposure time of 8 s and a frame time of 200 ms.

Grids for the two encapsulin variant samples **S5D** and **S7G** were prepared by applying 3 μL of purified encapsulin sample at a concentration of 2 mg/mL to holey carbon 2/2 C-flat grids. This was followed by plunge-freezing in liquid ethane using a FEI Vitribot Mark IV (95% humidity, blot force: 0, wait time: 30, temperature: 4 °C).

Cryo-EM movies of samples **S5D** and **S7G** were collected at University of New South Wales on an Arctica cryo-electron microscope (Thermo Fisher Scientific) operating at 200 kV equipped with a Falcon III EC detector. Movies were collected with a calibrated pixel size of 0.986 Å using Thermo Fisher EPU automated data collection software. The exposure time was 47.84 s with a 1.196 s frame time. Further details and parameters of cryo-EM data collection can be found in SI Tables S6.5 and S6.6.

All cryo-EM data processing was performed using cryoSPARC^65^ v2.15.0. After importing movies, beam-induced and local motion was corrected using Patch motion correction. CTF correction was carried out using the Patch CTF estimation job. Then, 200 particles were manually picked using the Manual picker followed by 2D classification with 10 classes. Good classes were then used as templates for template-based particle picking via the Template picker. Picks were visually inspected to find optimal Local power and NCC score values. Particles were extracted with a box size of 384 pixels followed by two rounds of 2D classification using 100 and then 50 classes and otherwise standard parameters. Particle stacks were further cleaned via 3D classification using the Heterogeneous Refinement job with 3 classes, icosahedral symmetry (I) enforced and a refinement box size of 256 voxels. As initial models, volumes based on the X-ray structure of the *T. maritima* encapsulin (PDB ID: 3DKT) generated in UCSF Chimera 1.13.1rc were used. Particles of the largest 3D class were then used to carry out 3D refinement using the Homogeneous refinement (New) (I symmetry enforced, refinement box size: 384 voxels) job followed by Local and then Global CTF refinement with standard parameters. This was repeated 3× followed by a final round of 3D refinement. Local resolution estimation of final maps was done in cryoSPARC with the FSC threshold set to 0.143. Further details about cryo-EM data processing can be found in SI Tables S6.5 and S6.6 and SI Figures S6.1, S6.2 and S6.3.

### Model building and refinement

Final cryoSPARC maps were opened in UCSF Chimera^66^ version 1.14. An initial model was placed into the density by manually aligning a monomer of the *T. maritima* encapsulin (PDB ID: 3DKT) to the map, followed by the fit_to_volume command to improve the rigid body fit into the density. Coordinates were then opened in Coot^67^ version 0.9-pre and manually refined against the map. Briefly, the refinement strategy consisted of an initial rigid body refinement into the map, followed by chain refinement, and finally iterative manual real space refinements along the length of the protein. During refinement, density for a flavin-based molecule was identified between residues W87 and R78. Because the exact flavin derivative could not be identified from the density alone, riboflavin (RBF) was placed in the density and refined. Following manual refinements in Coot, coordinates were further refined as monomers in Phenix^68^ version 1.18.2-3874 using phenix.real_space_refine with default parameters. Restraints for the riboflavin were generated using eLBOW^69^ in Phenix. Non-crystallographic symmetry (NCS) was identified in the cryo-EM map via phenix.map_symmetry, and NCS operators were then applied using phenix.apply_ncs to construct a complete model with icosahedral symmetry. This model was then refined using phenix.real_space_refine with NCS restraints, global minimization, and ADP refinements. Models were then validated using the Comprehensive Validation (phenix.validation_cryoem) tool^70^ in Phenix prior to submitting to the PDB. Models were assigned the following PDB identifiers: 7LII, 7LIJ, 7LIK, 7LIL, 7LIM, 7LIS, and 7LIT (SI Tables S6.5 and 6.6). Final cryo-EM maps were submitted to the EMDB with the following identifiers: 23379, 23380, 23381, 23382, 23383, 23384 and 23385.

### Pore size measurements

The HOLE package (2.2.005) was used for measuring the radius of the 5-fold encapsulin pores.^42^ The *T. maritima* encapsulin crystal structure 3DKT was used for measurements of the wild-type encapsulin,^28^ while measurements of cage variants were based on cryo-EM structures. The default file was used to determine atomic Van der Waals radii. Measurements were conducted with a cut-off radius of 20 Å to analyse the interior and exterior surfaces of the pore in addition to the channel.

### Synthesis and purification of LBT control peptide

Peptide synthesis was done on a Biotage Syro I Automated Parallel Peptide Synthesizer. Rink amide resin was treated with a solution of 40% piperidine in DMF (1.6 mL) for 4 min followed by a solution of 20% piperidine in DMF for 6 min. The resin was washed with DMF (4 × 2.5 mL). Fmoc-AA-OH (0.8 mL, 0.5 M), DIC (0.8 mL, 0.5 M), and Oxyma (0.8 mL, 0.55 M) were dispensed from stock solutions in DMF onto the resin. The sample was heated to 75 °C and mixed for 15 min. Upon completion, the resin was washed with DMF (4 × 2.5 mL). For capping, the resin was treated with a solution of 2.5% acetic anhydride and 5% DIPEA in DMF (1.6 mL) for 6 min. The resin was then washed with DMF (4 × 2.5 mL).

Resin containing synthesised peptide was cleaved with a cocktail consisting of 90:5:5 TFA:water:triisopropylsilane with agitation for 1 h. The volume of the resulting peptide solution was reduced under a stream of nitrogen before being triturated with diethyl ether. The solid crude peptide was centrifuged, and the pellet was dissolved in 50:50 water:acetonitrile. The crude peptide solution was purified by reverse-phase HPLC (Sunfire C18 semi-prep 10 × 250 mm; Solvent A: water + 0.1% TFA; Solvent B: acetonitrile + 0.1% TFA) with a 0-100% gradient over 20 min (peptide eluted at 14.1 min). Peptide fractions were combined and lyophilised to yield the pure LBT peptide (SI Section 5).

### Molecular dynamics simulations

#### System preparation

Encapsulin variants were prepared for simulation from their corresponding structures. For mutants **S1R, S1K, S3E, S5D, S6E, S7D, S7G**, the cryo-EM structures determined herein were used, while for the wild-type *T. maritima* encapsulin, the available crystal structure (PDB 3DKT)^28^ was used. The biological assembly of each structure was rebuilt, and a single five-fold pore was isolated. Two sets of simulation systems were then prepared using the AMBER20 suite of programs,^71^ one containing only enough Na^+^ to neutralize the protein (termed **Neutral**, used for pore size determination) and another containing 50 mM of TbCl_3_ to assess flux through the five-fold pores (termed **Flux**). Both sets of systems were solvated in cuboid box of TIP3P^72^ water with a minimum of 20 Å to the box edge. The protein was modeled with the AMBER ff14SB forcefield.^73^ For the **Neutral** simulation set, Na^+^ ions employed the Joung-Cheatham model^74^. For the **Flux** simulation set, the 12-6-4 ion model was used for both Tb^3+^ and Cl^−^.^75,76^ The 12-6-4 model has been demonstrated to provide good performance in its description of ion-water, ion-nucleotide and ion-protein interactions for divalent and trivalent ions.^75,77–79^ For simulations employing the 12-6-4 model, ParmEd was used to modify the *prmtop* topology file to add the C_4_ term to the Lennard-Jones matrix.

We note that the 50 mM TbCl_3_ concentration value was chosen in the **Flux** simulations to roughly match the total ionic strength of the buffer (100 mM HEPES) in the stopped-flow kinetics experiments. While this supplies an excess of Tb^3+^ relative to the kinetics experiments (max. 500 µM Tb^3+^ due to excessive background signal above this concentration), the comparisons between variants remain valid.

All simulations are summarized in SI Table 8.1, while the full simulation and analysis details can be found in SI Section 8.

#### Equilibration

Each system was then energy minimized with 5000 steps of steepest descent minimization. The minimized structures were then thermalized to 300 K over 2 ns with heavy atom restraints of 10 kcal/mol/A^2^ on all protein C_α_ atoms. Constant temperature was maintained using a Langevin thermostat with a coupling constant of 3.0 ps. Hydrogen-heavy atom bonds were constrained using the SHAKE algorithm^80^ so a timestep of 2 fs could be used. Long range electrostatics were calculated using PME^81,82^ with a real-space cutoff of 8 Å. All production simulations employed the pmemd.cuda GPU engine^82,83^ and the SPFP precision model.^84^ The heated systems were then simulated at constant temperature and box volume for 10 ns, using the same thermostat described above. As above, C_α_ restraints of 10 kcal/mol/A^2^ were employed to retain the structure of the encapsulin in the absence of the majority of the protein cage. Following simulation at constant temperature and box volume, each system was simulated at constant temperature and pressure for 10 ns. The Langevin thermostat described above was used to maintain the temperature at 300K. An isotropic Berendsen barostat^85^ was used to maintain the pressure at 1 bar, using a coupling constant of 2.0 ps. As above, C_α_ restraints of 10 kcal/mol/A^2^ were employed.

We note that simulations with TbCl_3_ exclusively on one side of the pentamer (ie. the experimental starting conditions during stopped-flow experiments) not attempted. Even without an ion gradient as a driving force, the flux of ions through the pore under these conditions is able to provide a suitable measure of pore permeability.

#### Production

Each **Neutral** system was then split into three replicates for production simulation, while each **Flux** systems was split into six replicates. Each replicate was assigned new velocities. Production runs were conducted for 500 ns per replicate for the **Neutral** systems and 400 ns per replicate for the **Flux** systems (see SI Table S8.1) at constant temperature (300 K) and pressure (1 bar) using the same thermostat and barostat detailed above. To ensure native-like motion of the pores while retaining the overall structure of the protein cage, a subset of protein C_α_ atoms had restraints applied. In the wild-type, residues 1-121 and 221-255 had C_α_ restraints of 10 kcal/mol/A^2^ applied. These residues correspond to the outer edges of the five-fold pore simulation assembly and comprise the primary beta sheet region and first two large helices of each subunit. Analogous residues in each mutant structure were also restrained. A visual representation of which areas of the structure had restraints applied is shown in SI Figure S8.3. Trajectories were saved every 100 ps.

#### Analysis

Analysis of all simulations was conducted using CPPTRAJ^86^ and the MDAnalysis python library.^87,88^ Simulations were visualized with VMD 1.9.3^89^ and Pymol 1.8.^90^ To assess ion flux through encapsulin pores, an analysis tool was developed based on the MDAnalysis library. In this approach, a cylinder was defined using proximal pore residues, with the central plane in the center of the pore. A flux event was determined to occur when an ion transited from the upper half of the cylinder to the bottom half and into bulk solution or vice versa. All molecular dynamics topologies and analysis tools used in this work can be found at: https://github.com/LauGroup/TmEnc_Flux

Statistical one-way ANOVA tests were conducted in GraphPad Prism using default settings.

Supplementary videos at 12 frames per second are included to illustrate the following:

1. An example simulation replicate run on the wild-type encapsulin.
2. The trajectory of two chloride anions through the five-fold pore of variants **S6E**.
3. Tb^3+^ ions bound to the wild-type encapsulin surface throughout a simulation replicate.

### Stopped-flow kinetic assays

Stopped-flow kinetic assays for monitoring luminescence turn-on were performed using a KinetAsyst stopped-flow spectrophotometer (TgK Scientific). A Hg/Xe lamp was used for excitation at 282 nm, while all emission transmitted through a 455 nm longpass filter was integrated for data analysis.

Encapsulin samples for stopped-flow were buffer exchanged and prepared at a monomer concentration of 8 µM in 100 mM HEPES-KOH, pH 7.6. TbCl_3_ solutions were prepared at a concentration of 500 µM in 1 mM HCl, with solutions for each analysis made immediately prior to measurement by dilution from a 50 mM TbCl_3_ stock solution in 1 mM HCl.

The temperature of the stopped-flow sample handling unit was set to 15 °C. Prior to each stopped-flow analysis, encapsulin and TbCl_3_ solutions were equilibrated in the sample handling unit for 10 min prior to analysis.

Each measurement was performed by injecting 50 µL of each solution into the sample chamber and recording luminescence for 2 s (8192 data points, 48 oversampling, 0.03 ms time constant). Rate constants were determined by averaging at least seven independent measurements and fitting the curve to a first-degree exponential (one-phase association in Prism 8 GraphPad), using a fitting window of 7-90 ms for encapsulin samples, and 3-50 ms for the LBT peptide. The quoted errors are the upper bound for the 95% confidence interval based on the model fitting (SI Table S7.6).

## Data availability

The sequence data, in GenBank format, for all the constructs created in this study are available on GitHub at https://github.com/LauGroup/TmEnc_Flux/tree/main/plasmid%20maps. Molecular dynamics topologies and input files are available on GitHub at https://github.com/LauGroup/TmEnc_Flux. All other data that support the findings of this study are available from the corresponding authors upon request.

## Acknowledgements

YHL acknowledges funding from the Australian Research Council (DE190100624) and the Westpac Foundation (WRF2020). TWG acknowledges support from the National Institutes of Health (R35GM133325). The authors acknowledge Assoc. Prof. Ronald Clarke for assistance with stopped-flow kinetics, Dr Anna Wang for assistance with interpreting kinetic data, Prof. Richard Payne for assistance with peptide synthesis, and Sydney Analytical for assistance with molecular characterization. Research reported in this publication was supported by the University of Michigan Cryo-EM Facility (U-M Cryo-EM). U-M Cryo-EM is grateful for support from the U-M Life Sciences Institute and the U-M Biosciences Initiative. The authors also acknowledge the facilities and the scientific and technical assistance of Microscopy Australia at the Australian Centre for Microscopy & Microanalysis at the University of Sydney and the Electron Microscopy Unit at the University of New South Wales, and the resources and services from the National Computational Infrastructure (NCI), which is supported by the Australian Government. The authors declare no competing interests.

