## Supplementary Information for "Pore structure controls stability and molecular flux in engineered protein cages"

### Table of contents — Page


| | | |
|---|---|---|
| 1. | **Materials** | **3** |
| 2. | **Molecular cloning**<br>**Table S2.1** Sequences for mutagenic primers used in cloning<br>**Table S2.2** Sequences of protein variants and LBT peptide | **4** |
| 3. | **Protein expression and purification**<br>**Figure S3.1** Example SDS-PAGE analysis of insoluble encapsulin variants<br>**Figure S3.2** Example FPLC chromatograph of initial crude size-exclusion chromatography of encapsulin variants<br>**Figure S3.3** Example FPLC chromatograph of ion-exchange chromatography of encapsulin variants<br>**Figure S3.4** Example FPLC chromatograph of final high-resolution size-exclusion chromatography of encapsulin variants, and analysis of purity by SDS-PAGE | **7** |
| 4. | **Characterization of purified encapsulin cage assemblies**<br>**Figure S4.1** TEM images of negatively stained wild type and encapsulin variants<br>**Figure S4.2** SEC chromatographs for SEC-MALS analysis of encapsulin variants<br>**Table S4.3** Molecular weight of purified encapsulin assemblies as determined by MALS, then hydrodynamic size and polydispersity index as determined by DLS<br>**Figure S4.4** The *T. maritima* encapsulin is a flavoprotein | **10** |
| 5. | **Characterization of synthetic lanthanide binding tag peptide**<br>**Figure S5.1** LCMS data for pure LBT peptide | **13** |
| 6. | **Cryo-electron microscopy (cryo-EM)**<br>**Figure S6.1** Cryo-EM data processing pipeline<br>**Figure S6.2** Additional Cryo-EM data<br>**Figure S6.3** Local resolution analysis of pore regions<br>**Figure S6.4** Cryo-EM data for embedded flavin<br>**Table S6.5** Cryo-EM data collection and refinement parameters for variants **S1K**, **S1R** and **S3E**<br>**Table S6.6** Cryo-EM data collection and refinement parameters for variants **S5D**, **S6E**, **S7D** and **S7G** | **14** |
| 7. | **Stopped-flow luminescence**<br>**Figure S7.1** Schematic and summary data from luminescence kinetic assay<br>**Figure S7.2** Example fitting of kinetic data to a single exponential curve<br>**Figure S7.3** Raw luminescence data curves<br>**Figure S7.4** Titration of Tb3+ against wild-type encapsulin tagged with *N*-terminal LBT<br>**Figure S7.5** Stopped-flow luminescence turn-on curves for LBT peptide and tagged wild-type encapsulin when varying initial input concentrations<br>**Table S7.6** Table of rate constants for $Tb^{3+}$ luminescence turn-on kinetics | **20** |
| 8. | **Molecular dynamics simulations**<br>**Table S8.1** Summary of all simulations conducted in this work<br>**Figure S8.2** Pore sizes calculated by the HOLE program during simulations<br>**Figure S8.3** Encapsulin pentamer showing areas with restraints applied<br>**Figure S8.4** Encapsulin pentamer showing cylinder for counting flux events | **24** |
| 9. | **Theoretical estimation of $Tb^{3+}$ flux** | **26** |
| 10. | **Supplementary references** | **27** |
| 11. | **Supplementary video files** (attachments)<br>**SI Video 1** An example simulation replicate run on the wild-type encapsulin<br>**SI Video 2** The trajectory of two chloride anions through the five-fold pore of variants S6E<br>**SI Video 3** $Tb^{3+}$ ions bound to the wild-type encapsulin surface throughout a simulation replicate | **–** |

#### 1. Materials

Oligonucleotides were synthesized by Sigma-Aldrich. PCR amplification was performed using Q5 High-Fidelity Master Mix obtained from New England BioLabs (NEB). PCR products were purified using the Monarch PCR and DNA Cleanup Kits (NEB).

SDS-PAGE was performed using Mini-PROTEAN TGX Stain-Free gels (Bio-Rad) with Tris-glycine-SDS running buffer. Gels were stained using Coomassie Brilliant Blue G-250. Unstained Protein Standard, Broad Range 10-200 kDa (NEB) was used as a ladder for all gels unless otherwise specified. Gel images were captured on a ChemiDoc MP Imaging System (BioRad).

Terbium(III) chloride hexahydrate was purchased from Sigma-Aldrich. All other buffer components and salts were purchased from either Sigma-Aldrich or Thermo Fisher Scientific.

#### 2. Molecular cloning

**Table S2.1 Sequences for mutagenic primers used in cloning.**

| Primer name | Sequence (5’ to 3’) |
|---|---|
| S1D fwd | GAGGCCGGGGACTATCCCTTGGAGAAA |
| S1D rev | AAGGGATAGTCCCCGGCCTCTTCTTTCAG |
| S2D fwd | GAAGAGGCCGATGACTATCCCTTGGAGAAACG |
| S2D rev | AAGGGATAGTCATCGGCCTCTTCTTTCAGG |
| S3D fwd | TGAAAGAAGAGGCCGATGACGATCCCTTGGAGAAAC |
| S3D rev | AAGGGATCGTCATCGGCCTCTTCTTTCAGGAAATTG |
| S4D fwd | GAAAGAAGATGACGATGACTTGGAGAAACGCGTAGAGG |
| S4D rev | TCTCCAAGTCATCGTCATCTTCTTTCAGGAAATTGAT |
| S5D fwd | TGAAAGAAGACGACTTGGAGAAACGCGTAGAGG |
| S5D rev | TTTCTCCAAGTCGTCTTCTTTCAGGAAATTGATC |
| S7D fwd | TCAATTTCCTGGACGATGAGAAACGCGTAGAG |
| S7D rev | ACGCGTTTCTCATCGTCCAGGAAATTGATCCAAC |
| S7G fwd | CAACTTCCTGGGCGGCGAGAAGAGAGTGGAAGAATGCC |
| S7G rev | TCTTCTCGCCGCCCAGGAAGTTGATCCATCTGTCCGTG |
| S1E fwd | AAGAGGCCGAGGAGTATCCCTTGGAGAAAC |
| S1E rev | CAAGGGATACTCCCCGGCCTCTTCTTTC |
| S2E fwd | GGCCGAGGAGGAGCCCTTGGAGAAACG |
| S2E rev | AGGGATACTCCTCGGCCTCTTCTTTCAG |
| S3E fwd | GGCCGAGGAGGAGCCCTTGGAGAAACG |
| S3E rev | AAGGGCTCCTCCTCGGCCTCTTCTTTCAG |
| S4E fwd | AAGAGGAGGAGGAGTTGGAGAAACGCGTAG |
| S4E rev | ACTCCTCCTCCTCTTCTTTCAGGAAATTGATC |
| S5E fwd | AAAGAAGAGGAGTTGGAGAAACGCGTAG |
| S5E rev | TCCAACTCCTCTTCTTTCAGGAAATTGATC |
| S6E fwd | ATTTCCTGAAAGAAGAGAAACGCGTAGAGGAG |
| S6E rev | GCGTTTCTCTTCTTTCAGGAAATTGATC |
| S1K fwd | GAGGCCGGGAAGTATCCCTTGGAGAAAC |
| S1K rev | AAGGGATACTTCCCGGCCTCTTCTTTC |
| S2K fwd | GAGGCCAAGAAGTATCCCTTGGAGAAACGC |
| S2K rev | CAAGGGATACTTCTTGGCCTCTTCTTTCAG |
| S3K fwd | GGCCAAGAAGAAGCCCTTGGAGAAACGC |
| S3K rev | AAGGGCTTCTTCTTGGCCTCTTCTTTCAG |
| S4K fwd | AAGAAGAAGAAGTTGGAGAAACGCGTAG |
| S4K rev | ACTTCTTCTTCTTTTCTTTCAGGAAATTGATC |
| S5K fwd | AGAAAAGAAGTTGGAGAAACGCGTAGAG |
| S5K rev | CCAACTTCTTTTCTTTCAGGAAATTGATCC |
| S6K fwd | TTTCCTGAAGAAGGAGAAACGCGTAGAG |
| S6K rev | TTTCTCCTTCTTCAGGAAATTGATCCAAC |
| S1R fwd | AGAGGCCGGGCGCTATCCCTTGGAGAAACG |
| S1R rev | CAAGGGATAGCGCCCGGCCTCTTCTTTC |
| S2R fwd | GAGGCCCGCCGCTATCCCTTGGAGAAAC |
| S2R rev | AGGGATAGCGGCGGGCCTCTTCTTTCAG |
| S3R fwd | GGCCCGCCGCCGCCCCTTGGAGAAACGC |
| S3R rev | AGGGGCGGCGGCGGGCCTCTTCTTTCAG |
| S4R fwd | TGAAAGAACGTCGACGTCGATTGGAGAAACGCGTAG |
| S4R rev | TTCTCCAATCGACGTCGACGTTCTTTCAGGAAATTGATC |
| S5R fwd | CTGAAAGAACGTCGATTGGAGAAACGCGTAGAGG |
| S5R rev | CGTTTCTCCAATCGACGTTCTTTCAGGAAATTGATCC |

| | |
|---|---|
| S6R fwd | TTTCCTGAAACGTGAGAAACGCGTAGAGG |
| S6R rev | CGCGTTTCTCACGTTTCAGGAAATTGATCC |
| **Primers for lanthanide binding tag addition** | |
| NLBT fwd | GCTCTTATTTTACTGAAATTCTGAAATTCGAAGCTTGGCTGTTT<br>TGGC |
| NLBT rev | CAAAAGAGCGCTTAAGGAACTCCCCAGATCTCCCCGCCAGCA<br>GTTCATC |
| Pore variant fwd | GGCGATGAACTGCTGGCGGGGAGATCTGGGGAGTTCCTTAAG<br>CGCTCTTTTG |
| Pore variant rev | GCCAAAACAGCCAAGCTTCGAATTTCAGAATTTCAGTAAAAT<br>AAGAGCCTCCGG |

**Table S2.2 Sequences of protein variants and LBT peptide.** The LBT sequence was appended to the *N*-terminus of encapsulin variants for kinetics experiments. Loop region is highlighted in yellow, with the six-membered loop in **bold**, and point mutations in **red**. Full plasmid maps are available on GitHub as GenBank files (https://github.com/LauGroup/TmEnc_Flux/tree/main/plasmid%20maps).

| Name | Sequence |
|---|---|
| Wild type *T. maritima* encapsulin | MEFLKRSFAPLTEKQWQEIDNRAREIFKTQLYGRKFVDVEGPYGWEY<br>AAHPLGEVEVLSDENEVVKWGLRKSLPLIELRATFTLDLWELDNLER<br>GKPNVDLSSLEETVRKVAEFEDEVIFRGCEKSGVKGLLSFEERKIECG<br>STPKDLLEAIVRALSIFSKDGIEGPYTLVINTDRWINFLKE**EAGHYP**LE<br>KRVEECLRGGKIITTPRIEDALVVSERGGDFKLILGQDLSIGYEDREKD<br>AVRLFITETFTFQVVNPEALILLKF |
| S1D | MEFLKRSFAPLTEKQWQEIDNRAREIFKTQLYGRKFVDVEGPYGWEY<br>AAHPLGEVEVLSDENEVVKWGLRKSLPLIELRATFTLDLWELDNLER<br>GKPNVDLSSLEETVRKVAEFEDEVIFRGCEKSGVKGLLSFEERKIECG<br>STPKDLLEAIVRALSIFSKDGIEGPYTLVINTDRWINFLKE**EAGDYP**LE<br>KRVEECLRGGKIITTPRIEDALVVSERGGDFKLILGQDLSIGYEDREKD<br>AVRLFITETFTFQVVNPEALILLKF |
| S1E | MEFLKRSFAPLTEKQWQEIDNRAREIFKTQLYGRKFVDVEGPYGWEY<br>AAHPLGEVEVLSDENEVVKWGLRKSLPLIELRATFTLDLWELDNLER<br>GKPNVDLSSLEETVRKVAEFEDEVIFRGCEKSGVKGLLSFEERKIECG<br>STPKDLLEAIVRALSIFSKDGIEGPYTLVINTDRWINFLKE**EAGEYP**LE<br>KRVEECLRGGKIITTPRIEDALVVSERGGDFKLILGQDLSIGYEDREKD<br>AVRLFITETFTFQVVNPEALILLKF |
| S1K | MEFLKRSFAPLTEKQWQEIDNRAREIFKTQLYGRKFVDVEGPYGWEY<br>AAHPLGEVEVLSDENEVVKWGLRKSLPLIELRATFTLDLWELDNLER<br>GKPNVDLSSLEETVRKVAEFEDEVIFRGCEKSGVKGLLSFEERKIECG<br>STPKDLLEAIVRALSIFSKDGIEGPYTLVINTDRWINFLKE**EAGKYP**LE<br>KRVEECLRGGKIITTPRIEDALVVSERGGDFKLILGQDLSIGYEDREKD<br>AVRLFITETFTFQVVNPEALILLKF |
| S1R | MEFLKRSFAPLTEKQWQEIDNRAREIFKTQLYGRKFVDVEGPYGWEY<br>AAHPLGEVEVLSDENEVVKWGLRKSLPLIELRATFTLDLWELDNLER<br>GKPNVDLSSLEETVRKVAEFEDEVIFRGCEKSGVKGLLSFEERKIECG<br>STPKDLLEAIVRALSIFSKDGIEGPYTLVINTDRWINFLKE**EAGRYP**LE<br>KRVEECLRGGKIITTPRIEDALVVSERGGDFKLILGQDLSIGYEDREKD<br>AVRLFITETFTFQVVNPEALILLKF |
| S2E | MEFLKRSFAPLTEKQWQEIDNRAREIFKTQLYGRKFVDVEGPYGWEY<br>AAHPLGEVEVLSDENEVVKWGLRKSLPLIELRATFTLDLWELDNLER<br>GKPNVDLSSLEETVRKVAEFEDEVIFRGCEKSGVKGLLSFEERKIECG<br>STPKDLLEAIVRALSIFSKDGIEGPYTLVINTDRWINFLKE**EAEEYP**LE |

| | |
|---|---|
| | KRVEECLRGGKIITTPRIEDALVVSERGGDFKLILGQDLSIGYEDREKD AVRLFITETFTFQVVNPEALILLKF |
| S3E | MEFLKRSFAPLTEKQWQEIDNRAREIFKTQLYGRKFVDVEGPYGWEY AAHPLGEVEVLSDENEVVKWGLRKSLPLIELRATFTLDLWELDNLER GKPNVDLSSLEETVRKVAEFEDEVIFRGCEKSGVKGLLSFEERKIECG STPKDLLEAIVRALSIFSKDGIEGPYTLVINTDRWINFLKE**EAEEEP**LE KRVEECLRGGKIITTPRIEDALVVSERGGDFKLILGQDLSIGYEDREKD AVRLFITETFTFQVVNPEALILLKF |
| S4D | MEFLKRSFAPLTEKQWQEIDNRAREIFKTQLYGRKFVDVEGPYGWEY AAHPLGEVEVLSDENEVVKWGLRKSLPLIELRATFTLDLWELDNLER GKPNVDLSSLEETVRKVAEFEDEVIFRGCEKSGVKGLLSFEERKIECG STPKDLLEAIVRALSIFSKDGIEGPYTLVINTDRWINFLK**EDDDD**LEKR VEECLRGGKIITTPRIEDALVVSERGGDFKLILGQDLSIGYEDREKDAV RLFITETFTFQVVNPEALILLKF |
| S5D | MEFLKRSFAPLTEKQWQEIDNRAREIFKTQLYGRKFVDVEGPYGWEY AAHPLGEVEVLSDENEVVKWGLRKSLPLIELRATFTLDLWELDNLER GKPNVDLSSLEETVRKVAEFEDEVIFRGCEKSGVKGLLSFEERKIECG STPKDLLEAIVRALSIFSKDGIEGPYTLVINTDRWINFLK**EDD**LEKRVE ECLRGGKIITTPRIEDALVVSERGGDFKLILGQDLSIGYEDREKDAVRL FITETFTFQVVNPEALILLKF |
| S5E | MEFLKRSFAPLTEKQWQEIDNRAREIFKTQLYGRKFVDVEGPYGWEY AAHPLGEVEVLSDENEVVKWGLRKSLPLIELRATFTLDLWELDNLER GKPNVDLSSLEETVRKVAEFEDEVIFRGCEKSGVKGLLSFEERKIECG STPKDLLEAIVRALSIFSKDGIEGPYTLVINTDRWINFLK**EEE**LEKRVE ECLRGGKIITTPRIEDALVVSERGGDFKLILGQDLSIGYEDREKDAVRL FITETFTFQVVNPEALILLKF |
| S6E | MEFLKRSFAPLTEKQWQEIDNRAREIFKTQLYGRKFVDVEGPYGWEY AAHPLGEVEVLSDENEVVKWGLRKSLPLIELRATFTLDLWELDNLER GKPNVDLSSLEETVRKVAEFEDEVIFRGCEKSGVKGLLSFEERKIECG STPKDLLEAIVRALSIFSKDGIEGPYTLVINTDRWINFLK**EE**KRVEECL RGGKIITTPRIEDALVVSERGGDFKLILGQDLSIGYEDREKDAVRLFITE TFTFQVVNPEALILLKF |
| S7D | MEFLKRSFAPLTEKQWQEIDNRAREIFKTQLYGRKFVDVEGPYGWEY AAHPLGEVEVLSDENEVVKWGLRKSLPLIELRATFTLDLWELDNLER GKPNVDLSSLEETVRKVAEFEDEVIFRGCEKSGVKGLLSFEERKIECG STPKDLLEAIVRALSIFSKDGIEGPYTLVINTDRWINFL**DD**EKRVEECL RGGKIITTPRIEDALVVSERGGDFKLILGQDLSIGYEDREKDAVRLFITE TFTFQVVNPEALILLKF |
| S7G | MEFLKRSFAPLTEKQWQEIDNRAREIFKTQLYGRKFVDVEGPYGWEY AAHPLGEVEVLSDENEVVKWGLRKSLPLIELRATFTLDLWELDNLER GKPNVDLSSLEETVRKVAEFEDEVIFRGCEKSGVKGLLSFEERKIECG STPKDLLEAIVRALSIFSKDGIEGPYTLVINTDRWINFL**GG**EKRVEECL RGGKIITTPRIEDALVVSERGGDFKLILGQDLSIGYEDREKDAVRLFITE TFTFQVVNPEALILLKF |
| Lanthanide binding tag (LBT) peptide | MGYIDTNNDGWYEGDELLAGRSG |

### 3. Protein expression and purification

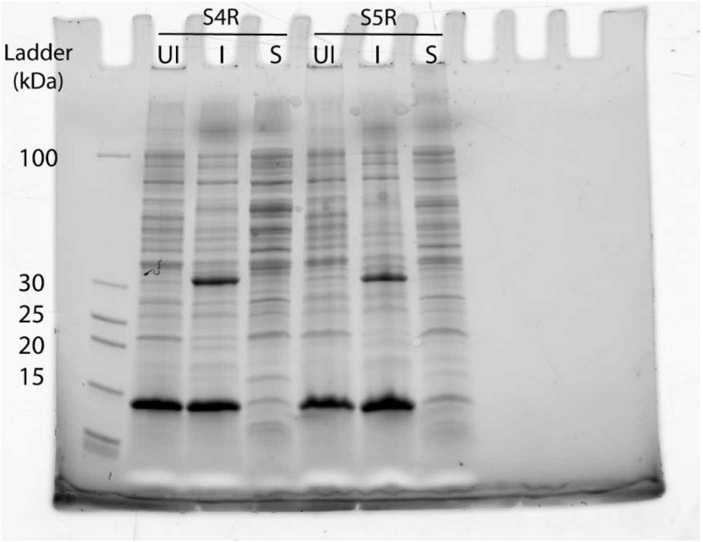


**Figure S3.1 Example SDS-PAGE analysis of insoluble encapsulin variants.** UI = un-induced control sample, I = insoluble proteins post-lysis, S = soluble proteins post-lysis. PageRuler unstained low range protein standard (ThermoFisher Scientific) was used as the ladder. Clear band for the encapsulin monomer (~32 kDa) is seen in the insoluble fraction for the variants **S4R** and **S5R**. Note, the dark band below 15 kDa is lysozyme added to aid cell lysis.

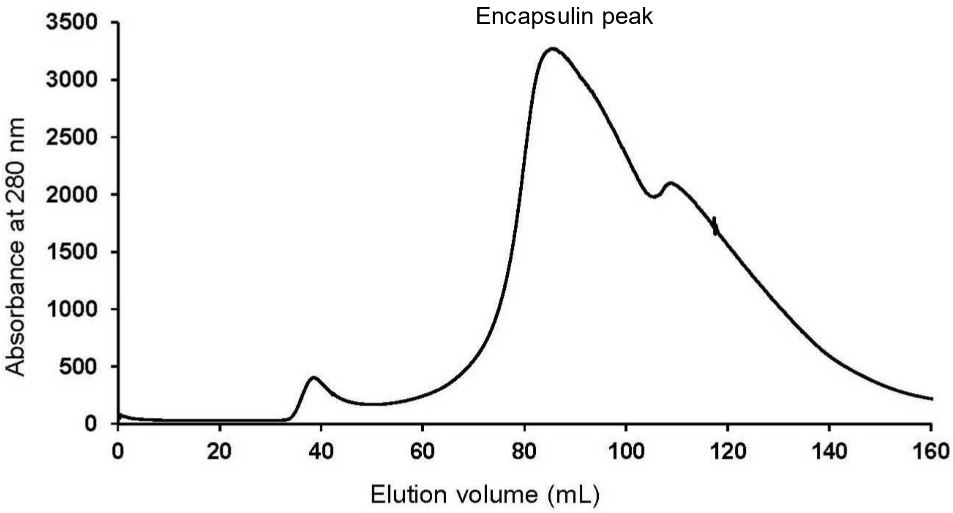


**Figure S3.2 Example FPLC chromatograph of initial crude size-exclusion chromatography of encapsulin variants.** Chromatography was performed on a Sephacryl S-500 16/60 column, with 20 mM Tris-HCl, 100 mM NaCl, pH 8 as the running buffer. Encapsulin was found to elute at ~65-110 mL.

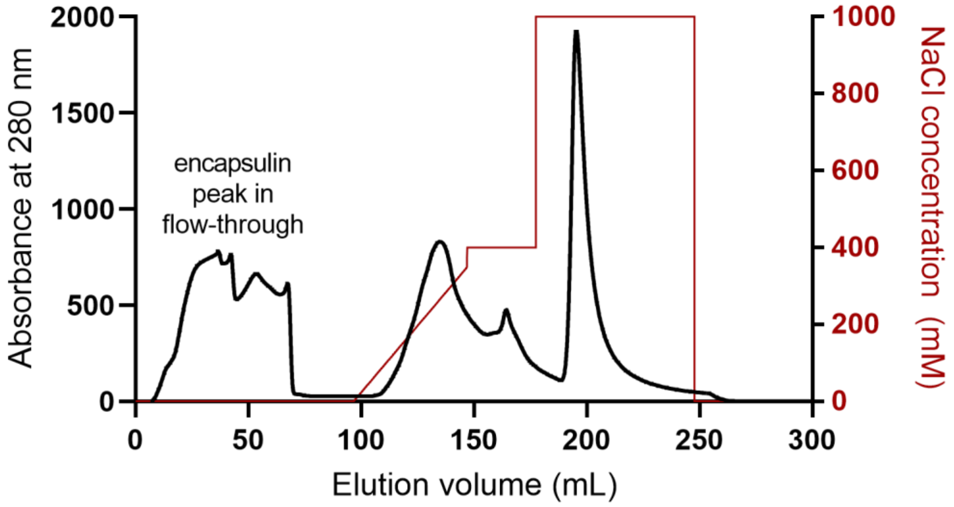


**Figure S3.3 Example FPLC chromatograph of ion-exchange chromatography of encapsulin variants.** Absorbance at 280 nm is shown by the black trace. Encapsulin peak is collected from the column flow-through void volume before applying a gradient. The red trace shows the NaCl concentration gradient used for the ion-exchange chromatography in the running buffer (20 mM Tris-HCl, pH 8).

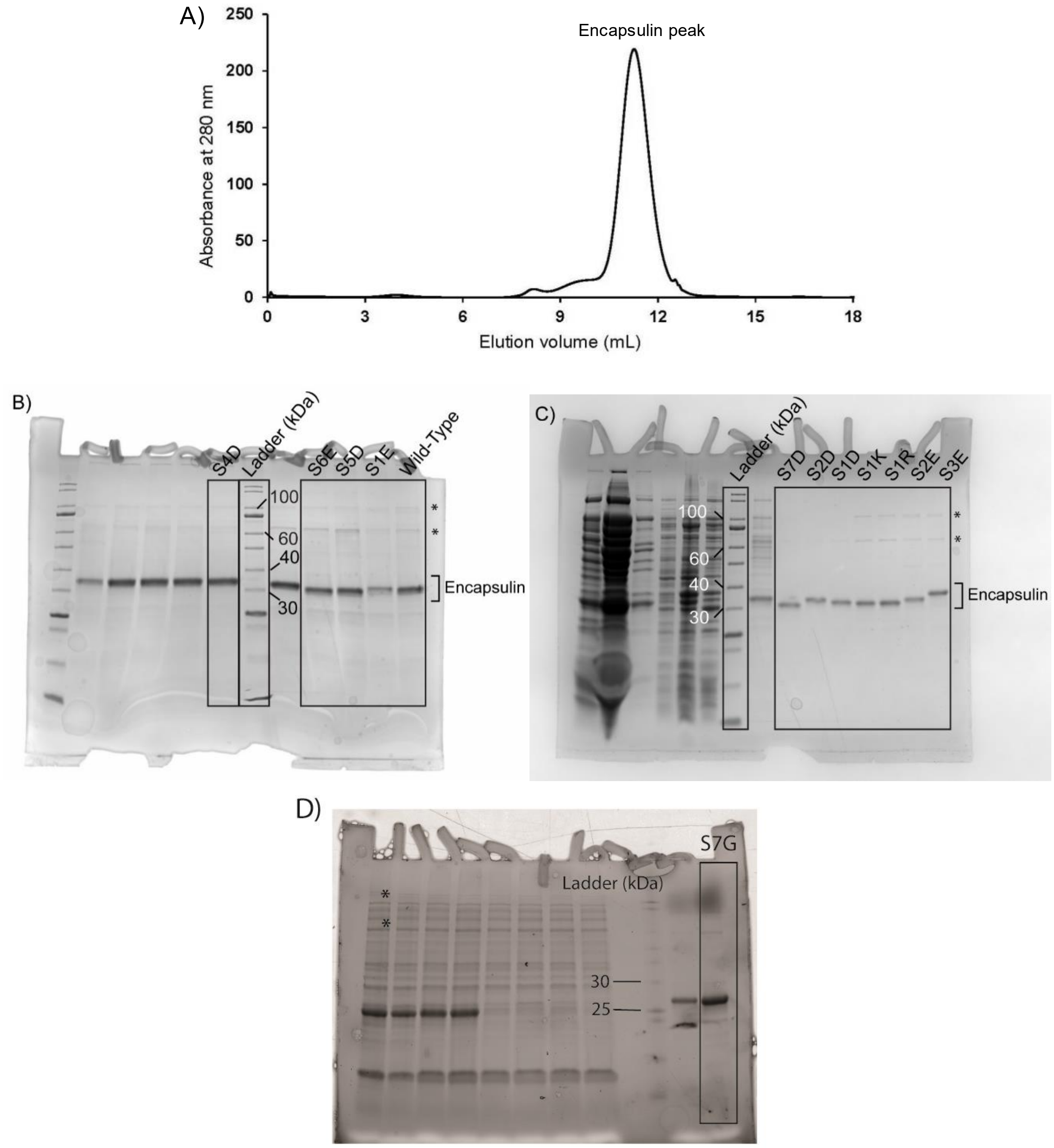


**Figure S3.4 A) Example FPLC chromatograph of final high-resolution size-exclusion chromatography of encapsulin variants.** The chromatograph shows the second round of size-exclusion chromatography using a Superose 6 10/300 GL column.

**B)-D) SDS-PAGE analysis of purified encapsulin variants.** The lanes corresponding to each variant are boxed and labelled above each lane. Gel B is the full uncropped gel from which the image in main text Figure 2aiv was taken.

These full unmodified gels are provided to explicitly show that no unjustified cropping or image editing has been carried out. The unlabelled lanes are not relevant to the data in this study.

Asterisks denote minor higher molecular weight bands in the gels that may correspond to encapsulin dimers (~66 kDa) and trimers (~99 kDa) that were not fully denatured due to high thermal stability.

#### 4. Characterization of purified encapsulin cage assemblies

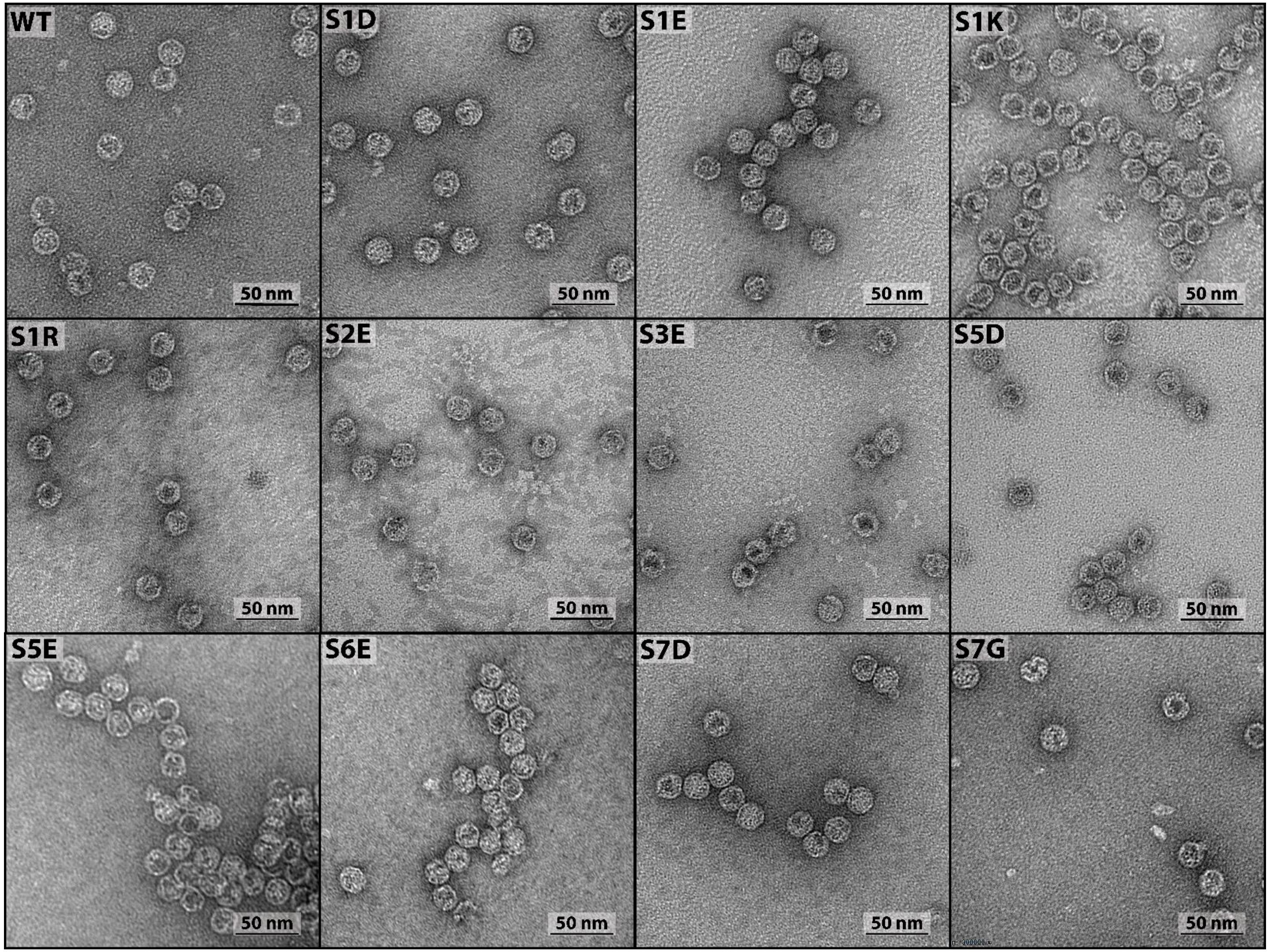


**Figure S4.1 TEM images of negatively stained wild type and encapsulin variants.**

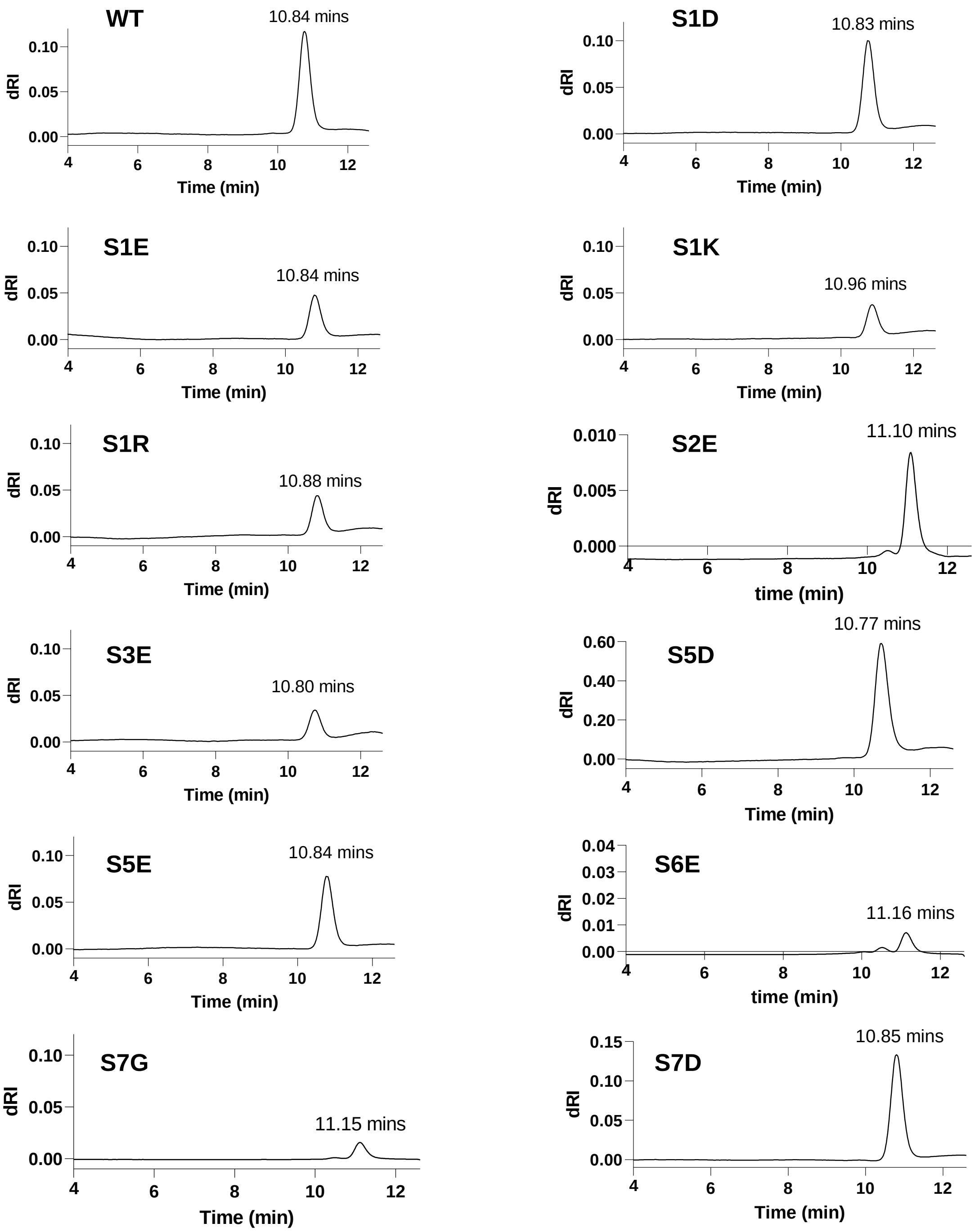


**Figure S4.2 SEC chromatographs for SEC-MALS analysis of encapsulin variants.** Chromatography run on a Bio SEC-5 column at a flow rate of 1 mL/min using PBS buffer, pH 7.4. The refractive index trace is shown.

**Table S4.3 Molecular weight of purified encapsulin assemblies as determined by MALS, then hydrodynamic size and polydispersity index as determined by DLS.**

| Encapsulin variant | Molecular weight (MDa) | Diameter (nm) | Polydispersity index ($M_w/M_n$) |
|---|---|---|---|
| WT | 2.17 ± 0.01 | 26.0 ± 2.0 | 1.06 ± 0.01 |
| S1D | 2.16 ± 0.01 | 24.5 ± 0.9 | 1.05 ± 0.01 |
| S1E | 2.51 ± 0.01 | 26.0 ± 2.0 | 1.03 ± 0.01 |
| S1K | 2.02 ± 0.02 | 25.0 ± 0.4 | 1.08 ± 0.01 |
| S1R | 2.14 ± 0.01 | 24.4 ± 0.3 | 1.07 ± 0.01 |
| S2E | 2.53 ± 0.01 | 25.8 ± 0.4 | 1.05 ± 0.01 |
| S3E | 2.11 ± 0.01 | 26.9 ± 0.7 | 1.05 ± 0.01 |
| S4D | 2.31 ± 0.01 | 24.4 ± 0.8 | 1.04 ± 0.01 |
| S5D | 2.70 ± 0.02 | 27.0 ± 0.2 | 1.06 ± 0.01 |
| S5E | 2.36 ± 0.01 | 25.8 ± 0.6 | 1.04 ± 0.01 |
| S6E | 2.07 ± 0.02 | 25.9 ± 0.6 | 1.01 ± 0.01 |
| S7D | 2.26 ± 0.01 | 24.7 ± 0.3 | 1.03 ± 0.01 |
| S7G | 2.41 ± 0.02 | 24.4 ± 0.3 | 1.01 ± 0.01 |

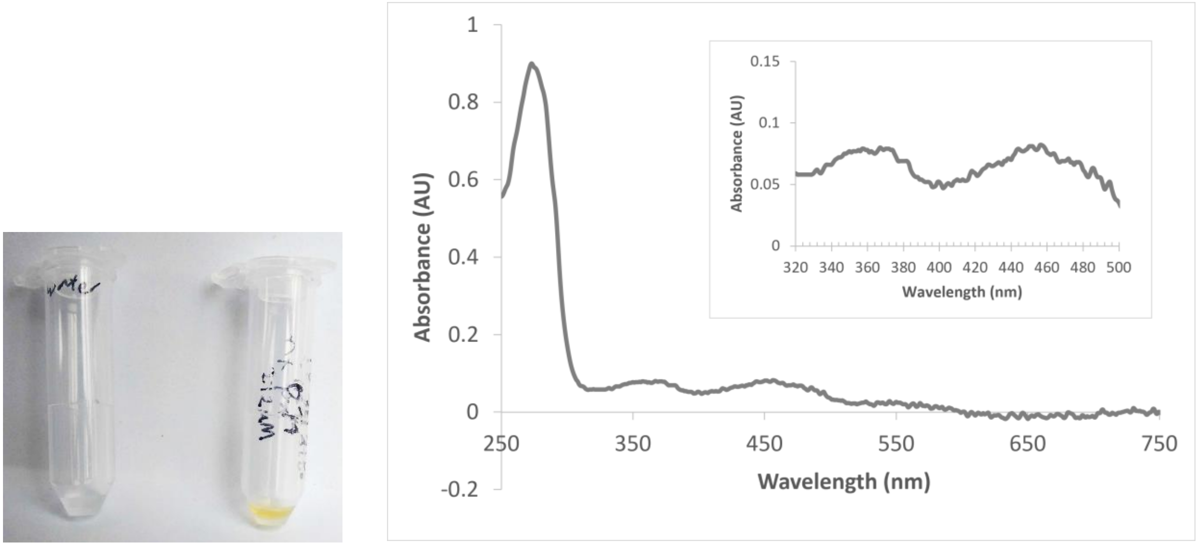


**Figure S4.4 The *T. maritima* encapsulin is a flavoprotein.**

**Left:** Pure concentrated encapsulin samples had a vibrant yellow colour (water shown in tube on the left for comparison).

**Right:** Clear absorbance maxima at approx. 450 and 370 nm corresponding to the presence of bound flavin (riboflavin, FMN or FAD). Inset shows zoom-in of the absorbance maxima. Cryo-EM data shows clear electron density corresponding to a flavin bound to each encapsulin monomer.

The presence of a flavin was confirmed in the literature during the preparation of this manuscript.[1]

#### 5. Characterization of synthetic lanthanide binding tag peptide

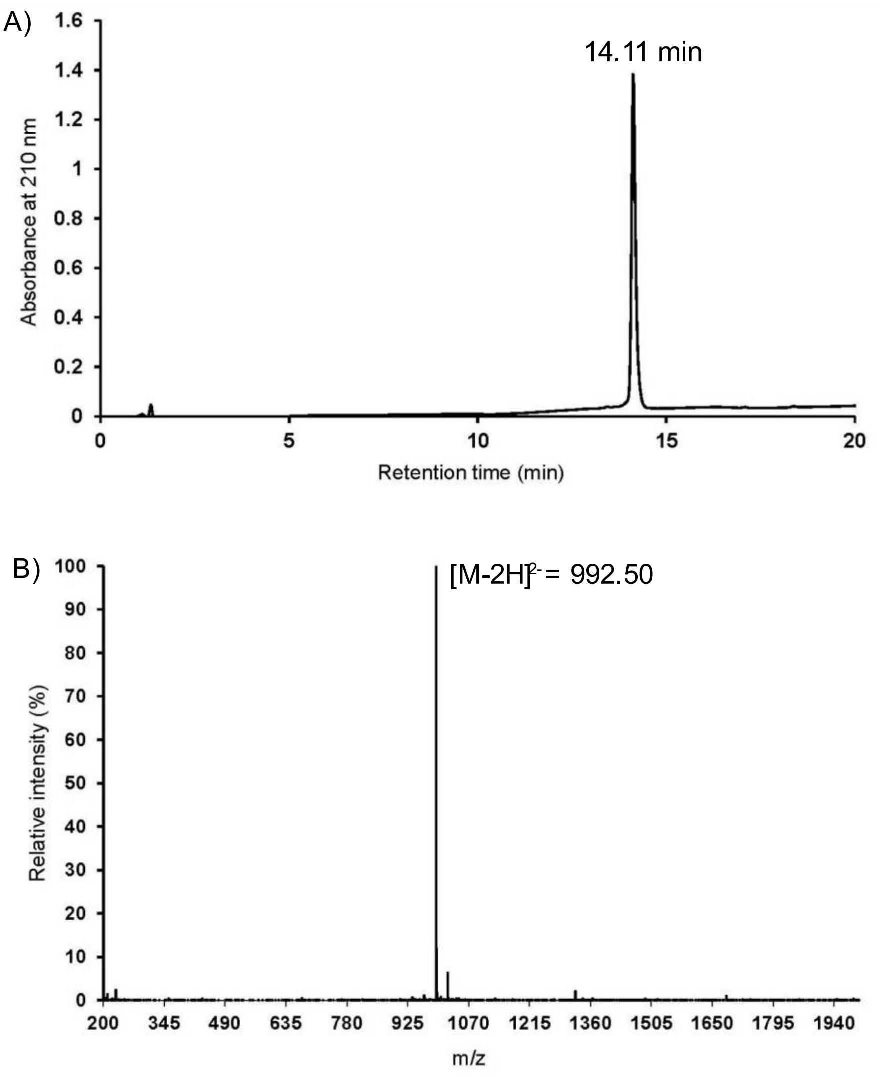


**Figure S5.1 LCMS data for pure LBT peptide.**

**A)** Analytical HPLC trace monitored at 210 nm, $R_t$ 14.11 min (0-100% acetonitrile over 20 min, 0.1% TFA).

**B)** Calculated mass: $[M+H]^+$ = 1988.06, $[M-2H]^{2-}$ = 992.52. Mass found (ESI-): 992.50 $[M-2H]^{2-}$.

#### 6. Cryo-electron microscopy

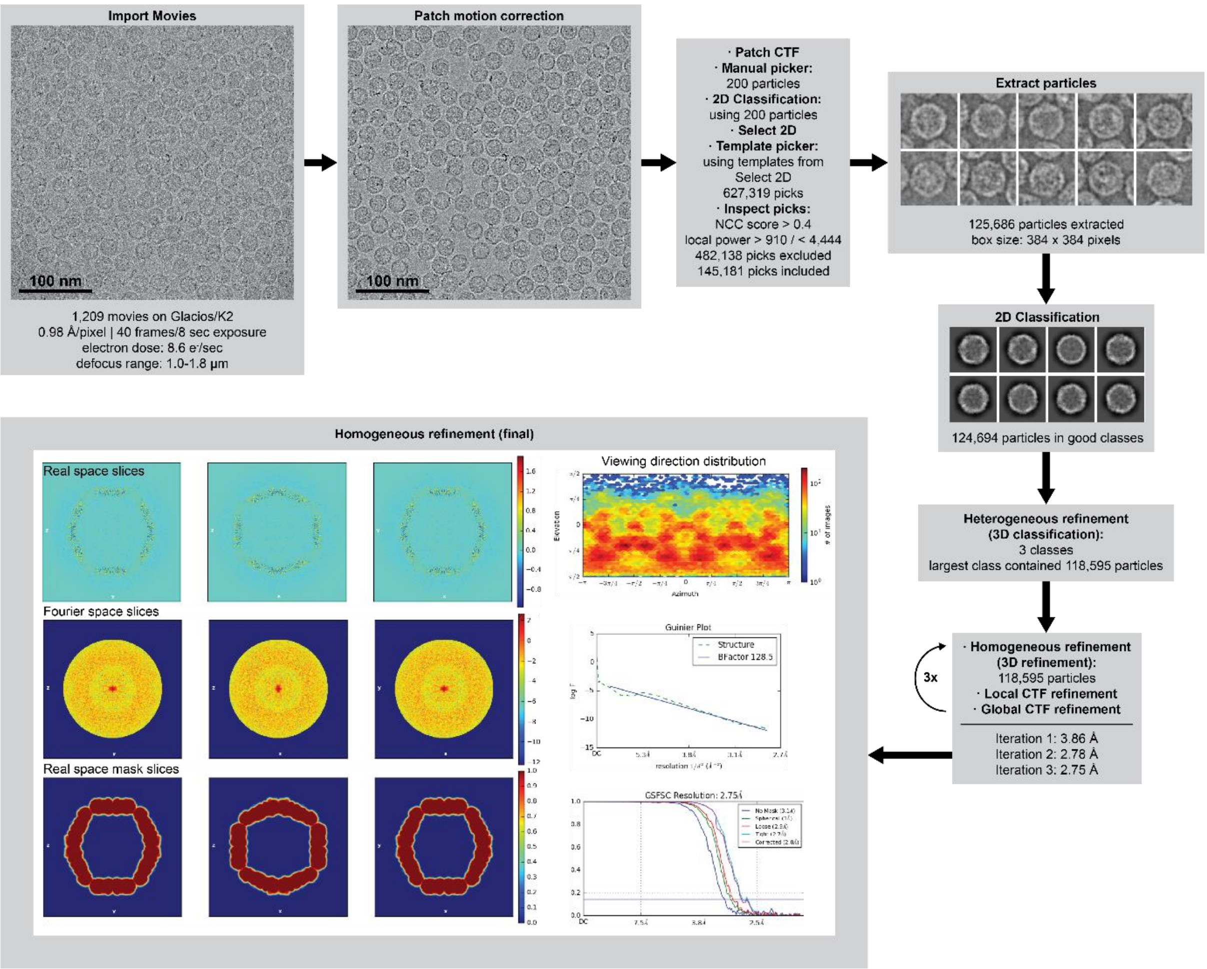


**Figure S6.1 Cryo-EM data processing pipeline.** An example cryo-EM data processing workflow is shown for variant **S6E**. All other variants were processed using the same workflow.

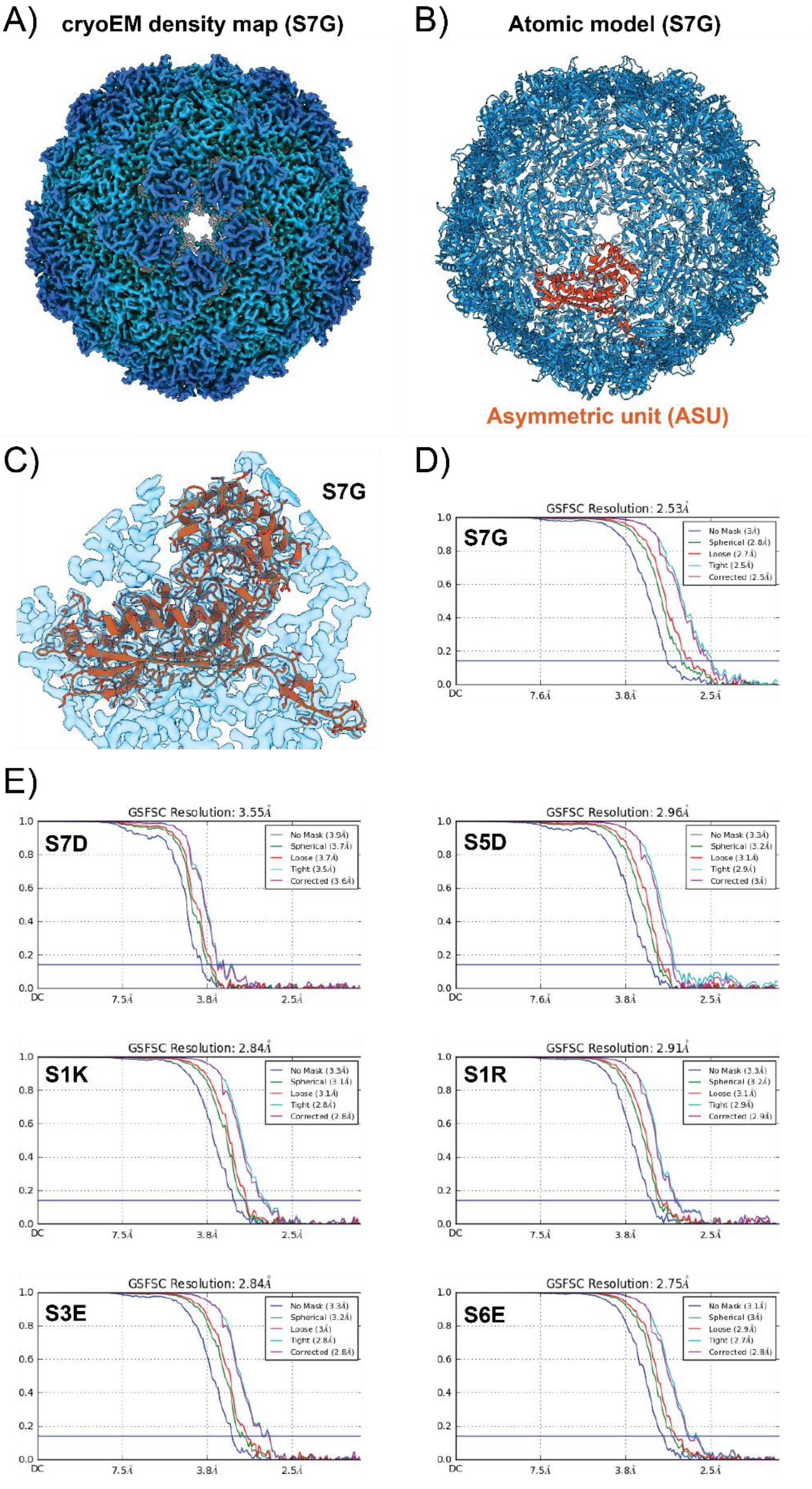


**Figure S6.2 Additional cryo-EM data. A)** Exterior view down the 5-fold symmetry axis of the cryo-EM map of variant **S7G**. **B)** Atomic model of variant **S7G** and highlighted asymmetric unit. **C)** Fit of a single **S7G** protomer into its cryo-EM density. **D)** Gold-standard FSC curves for variant **S7G** based on 0.143 threshold. **E)** Gold-standard FSC curves of the remaining variants based on 0.143 threshold.

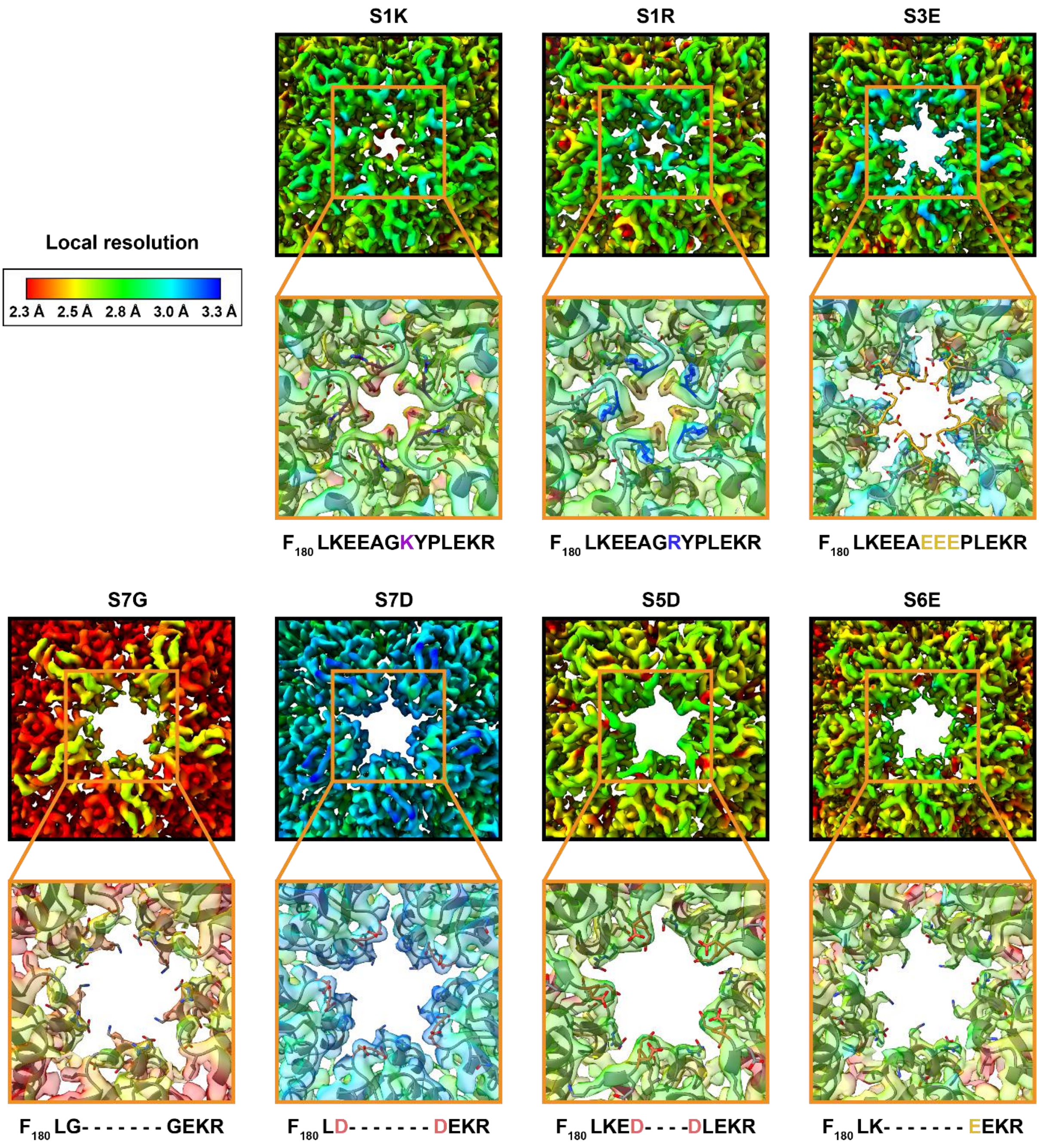


**Figure S6.3 Local resolution estimation for all variant cryo-EM maps.** For each variant, cryo-EM density coloured by local resolution is shown on top, while a zoom-in of the atomic model fitted into the cryo-EM density is shown below.

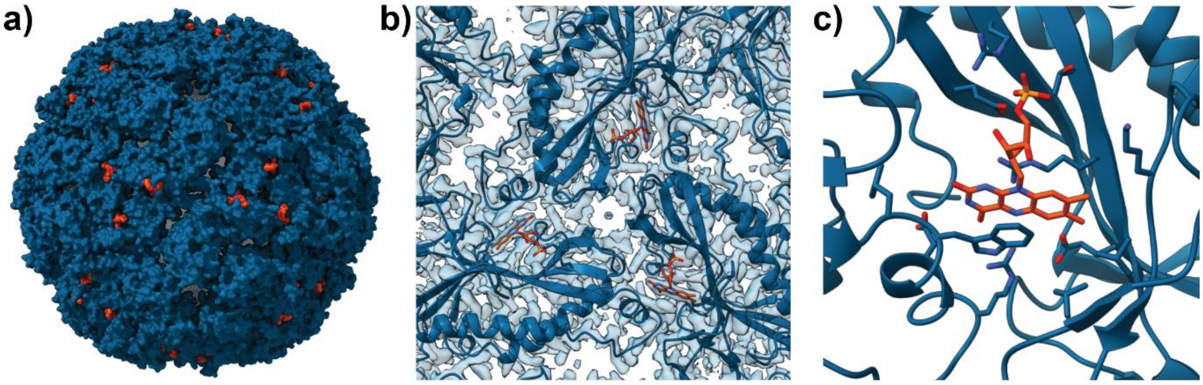


**Figure S6.4 Cryo-EM data for embedded flavin. a)** Surface view of the shell of mutant **S6E** with flavin co-factors arranged around the 3-fold pore highlighted in orange. **b)** Cryo-EM density and atomic model of mutant **S6E** around the 3-fold pore shown in cartoon representation. Flavin residues shown as stick models and colored orange. **c)** Flavin (orange) binding site. All residues within 5 Å are shown as stick models.

**Table S6.4 Cryo-EM data collection parameters and refinement statistics for variants S1K, S1R and S3E.**

| **Data Collection and Processing** | **S1K** | **S1R** | **S3E** |
|---|---|---|---|
| EMDB Accession | EMD-23380 | EMD-23381 | EMD-23382 |
| Electron Microscope | Glacios | Glacios | Glacios |
| Voltage (kV) | 200 | 200 | 200 |
| Total Dose ($e^-/Å^2$) | 62 | 62 | 69 |
| Defocus range (um) | -1.3 to -1.8 | -1.3 to -1.8 | -1.0 to -1.8 |
| Voltage (kV) | 200 | 200 | 200 |
| Particles | 58162 | 75829 | 39859 |
| Resolution (Å) | 2.84 | 2.91 | 2.84 |
| Map sharpening B-factor ($Å^2$) | -128.0 | -138.2 | -120.5 |
| Symmetry for map | I | I | I |
| Number of Movies | 911 | 1362 | 799 |
| **Structure Refinement (ASU)** | | | |
| PDB Accession | 7LIJ | 7LIK | 7LIL |
| Structure B factor (mean) | 22.03 | 27.84 | 23.67 |
| Ligand B factor (mean) | 17.30 | 25.58 | 13.95 |
| Mean CC for Protein (Mask) | 0.85 | 0.86 | 0.86 |
| Mean CC for Ligands | 0.74 | 0.71 | 0.75 |
| Chains in ASU | 1 | 1 | 1 |
| Bond lengths (Å) | 0.006 | 0.007 | 0.006 |
| Bond Angles (°) | 1.089 | 1.145 | 1.044 |
| **Ramachandran Statistics** | | | |
| Favoured % | 94.30 | 93.16 | 94.66 |
| Allowed % | 5.70 | 6.84 | 5.34 |
| Disallowed % | 0.00 | 0.00 | 0.00 |
| Rotamer Outliers (%) | 3.85 | 2.99 | 5.56 |
| **MolProbity Score** | 1.92 | 1.78 | 1.88 |
| **Clashscore** | 2.98 | 2.06 | 1.84 |

**Table S6.5 Cryo-EM data collection parameters and refinement statistics for variants S5D, S6E, S7D and S7G.**

| **Data Collection and Processing** | **S5D** | **S6E** | **S7D** | **S7G** |
|---|---|---|---|---|
| EMDB Accession | EMD-23384 | EMD-23383 | EMD-23379 | EMD-23385 |
| Electron Microscope | Arctica | Glacios | Glacios | Arctica |
| Voltage (kV) | 200 | 200 | 200 | 200 |
| Total Dose ($e^-$/Å$^2$) | 43 | 69 | 69 | 43 |
| Defocus range (um) | -1.0 to -2.5 | -1.0 to -1.8 | -1.0 to -1.8 | -1.0 to -2.5 |
| Voltage (kV) | 200 | 200 | 200 | 200 |
| Particles | 33416 | 118595 | 21501 | 90905 |
| Resolution (Å) | 2.96 | 2.75 | 3.55 | 2.53 |
| Map sharpening B-factor (Å$^2$) | -163.9 | -128.5 | -151.0 | -141.6 |
| Symmetry for map | I | I | I | I |
| Number of Movies | 623 | 1209 | 609 | 1301 |
| **Structure Refinement (ASU)** | | | | |
| PDB Accession | 7LIS | 7LIM | 7LII | 7LIT |
| Structure B factor (mean) | 51.86 | 37.36 | 41.41 | 61.68 |
| Ligand B factor (mean) | 52.70 | 27.82 | 40.47 | 56.33 |
| Mean CC for Protein (Mask) | 0.88 | 0.84 | 0.83 | 0.85 |
| Mean CC for Ligands | 0.74 | 0.78 | 0.75 | 0.78 |
| Chains in ASU | 1 | 1 | 1 | 1 |
| Bond lengths (Å) | 0.008 | 0.006 | 0.007 | 0.008 |
| Bond Angles (°) | 0.706 | 1.252 | 1.087 | 1.036 |
| **Ramachandran Statistics** | | | | |
| Favoured % | 96.53 | 96.48 | 90.23 | 96.09 |
| Allowed % | 3.47 | 3.52 | 9.77 | 3.91 |
| Disallowed % | 0.00 | 0.00 | 0.00 | 0.00 |
| Rotamer Outliers (%) | 3.45 | 3.49 | 0.87 | 0.44 |
| **MolProbity Score** | 1.62 | 1.83 | 1.77 | 1.14 |
| **Clashscore** | 2.09 | 3.99 | 4.48 | 1.42 |

Electrostatic maps shown in the manuscript were generated in PyMol 2.4.0 using the APBS Electrostatic plugin v.1.5.

### 7. Stopped-flow luminescence

The stopped-flow assay[2] involves genetically fusing a lanthanide binding tag (LBT) reported by Imperiali and co-workers to the *N*-terminus of the encapsulin gene,[3] thus resulting in 60 copies of the LBT displayed within the interior of every encapsulin protein cage. Upon binding to $Tb^{3+}$, energy transfer from a tryptophan in the LBT sequence to the lanthanide ion results in turn-on luminescence that can be measured using a stopped-flow apparatus and fitted to a one-phase association model. If the luminescent complex formation is extremely rapid, the rate constant of turn-on luminescence should reflect the kinetics of diffusion into the cage as the rate-determining step. However, our data suggests that diffusion is in fact faster than luminescent complex formation.

**Additional details of experimental protocol:** We observed a gradual reduction in luminescence signal with the age of prepared $TbCl_3$ solutions, due to precipitation of hydroxides when in buffered solution. To ascertain reproducibility, $TbCl_3$ stocks were prepared in 1 mM HCl, and diluted working solutions were freshly prepared prior to each experiment.

We also noted that Tris was an unsuitable buffer for kinetic experiments with $Tb^{3+}$, due to its ability to chelate the metal ion.

We ran negative control experiments by combining $TbCl_3$ + HEPES-KOH buffer, 1 mM HCl + LBT peptide, 1 mM HCl + LBT-tagged WT, or $TbCl_3$ + untagged WT, all of which showed no change in luminescence over the 2 s length of the experiment.

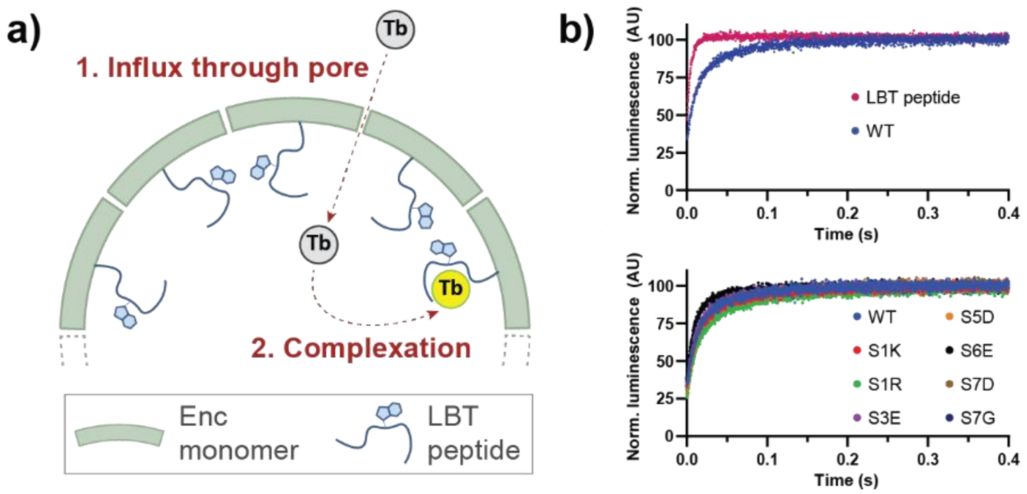


**Figure S7.1. a)** Schematic of kinetic assay, where a lanthanide binding tag (LBT) peptide is internally fused to each copy of the encapsulin, such that luminescence turn-on mechanism involves $Tb^{3+}$ diffusion through pores into the cage followed by complexation.[2,3] **b)** Top: Luminescence kinetic curves showing that the LBT peptide has a slower rate of turn-on when encapsulated. Bottom: Cage variants all show relatively similar rates of luminescence turn-on.

The fitting windows were chosen to capture the data that best fit a single exponential curve. Fitting the entire 2 s dataset as per *Williams et al*. gave a poorer fit,[2] indicating that the sample first-order approximation is not a complete model for the process (Figure S7.2). Nevertheless, while the 2 s fit gave slightly different rate constants, the relative trends between the variants remained similar.

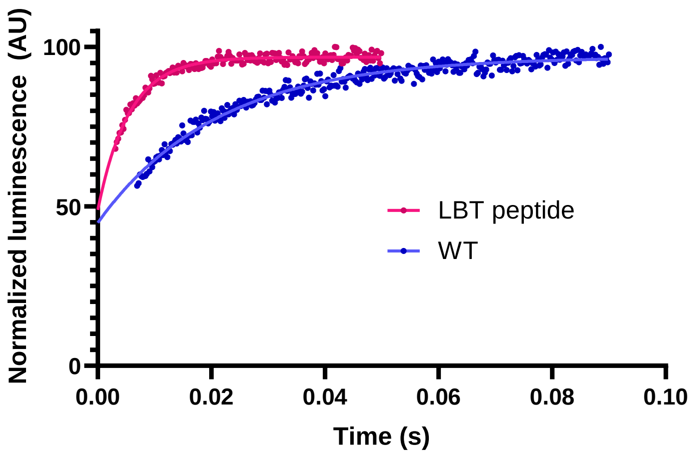


**Figure S7.2 Example fitting of kinetic data to a single exponential curve.** Data points within the fitting windows for the LBT peptide and the tagged wild-type encapsulin, corresponding to the data presented in Figure S7.1.

We also noted that the final luminescence intensity for the LBT peptide was significantly higher than that of all the encapsulated samples at the same concentration (Figure S7.3). This result may suggest that some form of crowding, self-quenching, inner-filter effect, or interior surface interaction is occurring within the encapsulin interior, causing the drop in luminescence intensity.

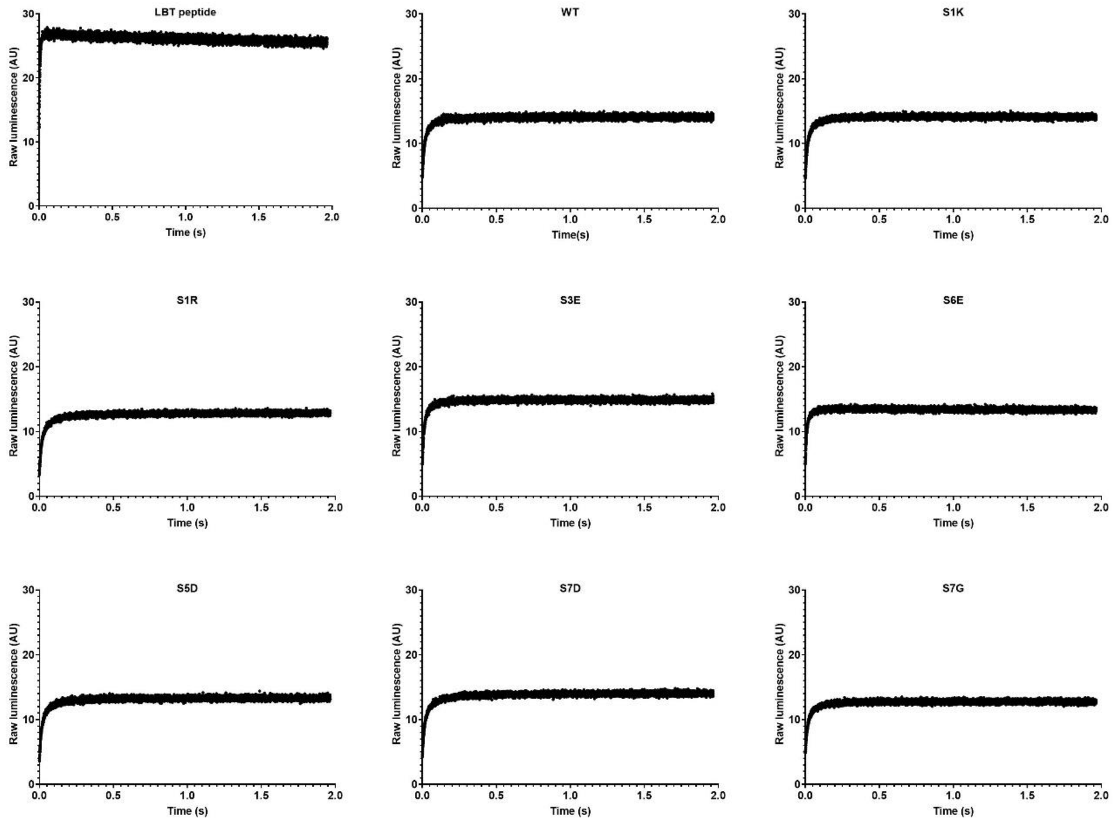

**Figure S7.3 Raw luminescence data curves.** Data corresponds to the bar graphs shown in Figure S7.1b.

A binding affinity titration was performed to verify that $Tb^{3+}$ was being complexed by the encapsulated LBT peptides as expected (Figure S7.4). Experiments were performed in technical triplicate. There were non-specific signal issues observed at higher $Tb^{3+}$ concentrations, attributed to background $Tb^{3+}$ luminescence.

Using a one-site binding model (total and non-specific, Prism 8 Graphpad), the binding affinity of the wild-type encapsulated LBT was approximated to be 660±40 nM, which is significantly weaker than the expected literature value of 57±4 nM for the LBT peptide.[3] This observation could potentially be one additional factor accounting for the slower on-rate observed in encapsulated samples.

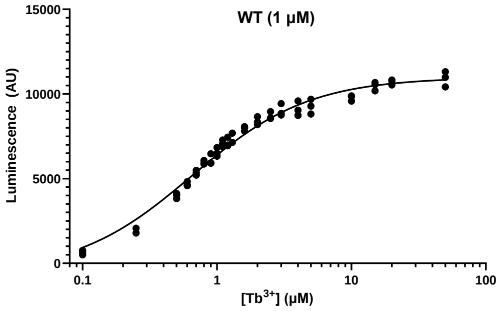


**Figure S7.4 Titration of $Tb^{3+}$ against wild-type encapsulin tagged with *N*-terminal LBT.**

Stopped-flow kinetics experiments were also run with varying $Tb^{3+}$ and LBT concentrations to determine their influence on the exponential fit constant (with implications for rate law).

Unusually for the free LBT peptide, the exponential constant did not vary substantially with changes in the initial concentration of either $Tb^{3+}$ or LBT peptide within the range tested (Figure S7.5) One possible interpretation is that an initial fast chelation event occurs between $Tb^{3+}$ and LBT peptide, followed by a rate-determining step such as a conformational change that is first order only with respect to the initial $Tb^{3+}$-LBT chelation complex.

In contrast, the tagged wild-type encapsulin did show a non-linear dependence on initial $Tb^{3+}$ concentration. This result indicates that the mechanistic processes governing luminescence turn-on kinetics are indeed different between free and encapsulated LBT peptide.

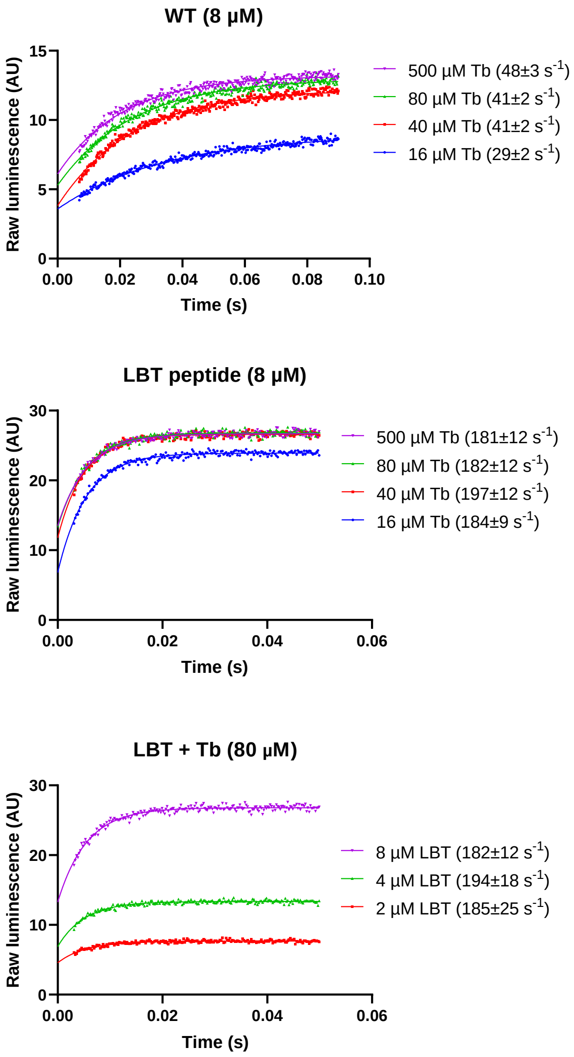


**Figure S7.5 Stopped-flow luminescence turn-on curves for LBT peptide and tagged wild-type encapsulin when varying initial input concentrations.**

| Cage variant | Rate constant ($s^{-1}$) |
|---|---|
| Free LBT peptide | 180 ± 10 |
| Wild-type (WT) | 48 ± 3 |
| S1K | 50 ± 2 |
| S1R | 44 ± 2 |
| S3E | 55 ± 3 |
| S5D | 43 ± 2 |
| S6E | 70 ± 4 |
| S7D | 44 ± 2 |
| S7G | 46 ± 2 |

**Table S7.6 Numerical values for $Tb^{3+}$ turn-on rate constants in kinetics experiments.** Corresponds to graph in main text Figure 4c.

### 8. Molecular dynamics simulations

**Table S8.1 Summary of all simulations conducted in this work.**

| Encapsulin | Simulation Type | Replicates | Replicate length | Total time | [$TbCl_3$] |
|---|---|---|---|---|---|
| **WT** | Neutral | 3 | 500 ns | 1.5 µs | - |
| **WT** | Flux | 6 | 400 ns | 2.4 µs | 50 mM |
| **S1K** | Neutral | 3 | 500 ns | 1.5 µs | - |
| **S1K** | Flux | 6 | 400 ns | 2.4 µs | 50 mM |
| **S1R** | Neutral | 3 | 500 ns | 1.5 µs | - |
| **S1R** | Flux | 6 | 400 ns | 2.4 µs | 50 mM |
| **S3E** | Neutral | 3 | 500 ns | 1.5 µs | - |
| **S3E** | Flux | 6 | 400 ns | 2.4 µs | 50 mM |
| **S5D** | Neutral | 3 | 500 ns | 1.5 µs | - |
| **S5D** | Flux | 6 | 400 ns | 2.4 µs | 50 mM |
| **S6E** | Neutral | 3 | 500 ns | 1.5 µs | - |
| **S6E** | Flux | 6 | 400 ns | 2.4 µs | 50 mM |
| **S7D** | Neutral | 3 | 500 ns | 1.5 µs | - |
| **S7D** | Flux | 6 | 400 ns | 2.4 µs | 50 mM |
| **S7G** | Neutral | 3 | 500 ns | 1.5 µs | - |
| **S7G** | Flux | 6 | 400 ns | 2.4 µs | 50 mM |

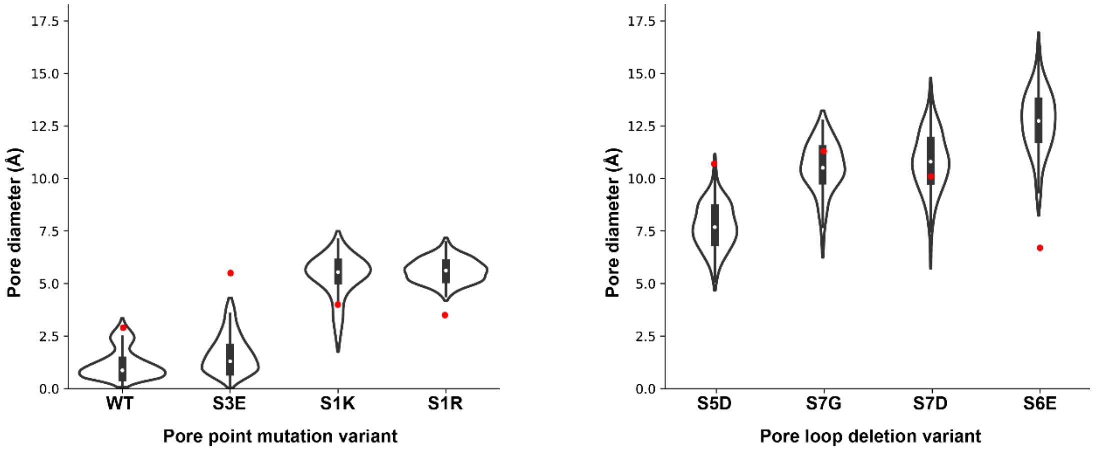


**Figure S8.2 Pore sizes calculated by the HOLE program during molecular dynamics simulations show that loop deletions possess greater pore flexibility.** The red dot represents the pore size based on the static cryo-EM structures. The violin plots show the distribution of pore sizes during the simulation window, with the box representing the interquartile range and the central line showing 1.5× the interquartile range. Point mutants and wild-type are displayed on the left plot, while the loop deletions are displayed on the right plot.

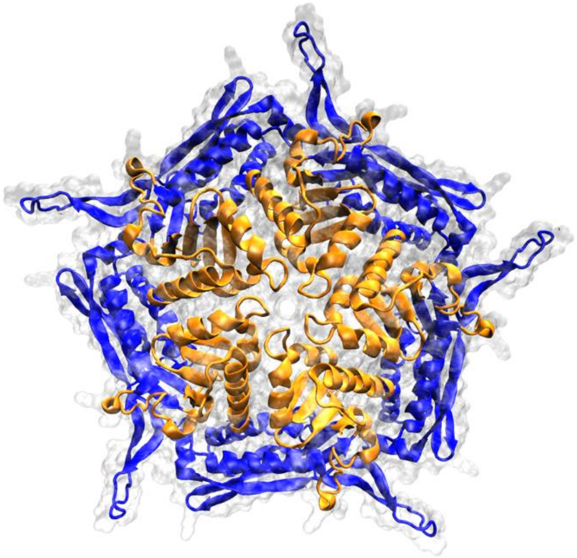

**Figure S8.3** Wild-type encapsulin pentamer structure showing the areas with $C_\alpha$ restraints applied (blue) and no restraints applied (orange).

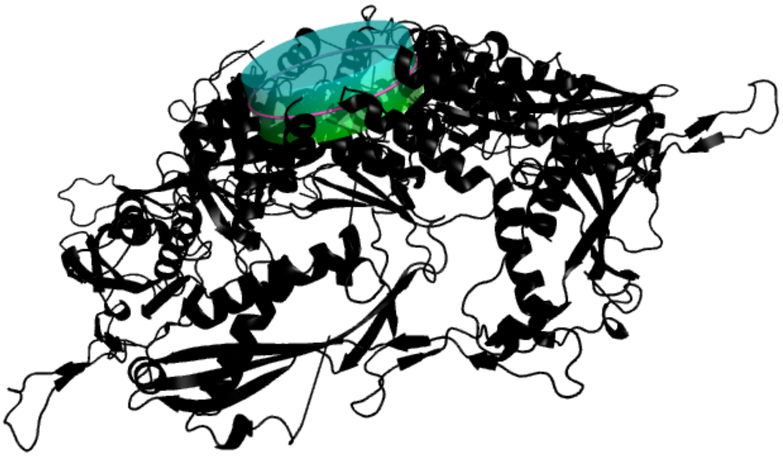

**Figure S8.4** Encapsulin mutant **S3E** pentamer structure showing the cylinder used for counting ion flux events, with upper (blue) and lower (green) sections divided at the center (pink).

#### 9. Theoretical estimation of $Tb^{3+}$ flux

Two models were used to estimate the rates of $Tb^{3+}$ diffusion into the wild-type *T. maritima* encapsulin compartment: one based on Berg's model of diffusion to $N$ disk-like adsorbers on the surface of a sphere,[4] and one based on Hinzpeter's model of carboxysome shell permeability.[5] Both models assume that the pores are inflexible and do not interact with $Tb^{3+}$, and that the diffusion coefficient of $Tb^{3+}$ at all times is equal to its diffusion coefficient in water. Neither model provides a complete description of the factors affecting diffusion into the encapsulin compartment. However, this estimation can provide a rough timescale for $Tb^{3+}$ diffusion into the encapsulin compartment for comparison to the stopped-flow experiments.

In Berg's model, the porous encapsulin shell is reduced to a reflecting sphere with disk-like adsorbers located on its surface in place of pores. In this model of diffusion, depletion of diffusing species close to the surface of the sphere is the main factor affecting diffusion. This model can be applied to the diffusion of $Tb^{3+}$ through the encapsulin under the assumption that the time taken for $Tb^{3+}$ to traverse the pore length is negligible. Using this model, the diffusive current is

$$I = \frac{4\pi D a C_0}{1 + \frac{\pi a}{N s}}$$

where $I$ is diffusive current in mol $s^{-1}$, $a$ is the radius of the encapsulin shell in m, $C_0$ is the initial $Tb^{3+}$ outside of the shell, $N$ is the number of pores, and $s$ is the pore radius in m. Assuming an encapsulin radius of 12 nm (the approximate radius for the *T. maritima* encapsulin), 62 pores with a radius of 1 Å (the number and approximate radius of pores in the *T. maritima* encapsulin shell), and an initial [$Tb^{3+}$] of 250 µM (the concentration used in the stopped-flow experiments), the diffusive current is

$$I = \frac{4 \cdot \pi \cdot (6 \cdot 10^{-10}\ m^2 s^{-1}) \cdot (1.2 \cdot 10^{-8}\ m) \cdot (2.5 \cdot 10^{-1}\ mol\ m^{-3})}{1 + \frac{\pi \cdot (1.2 \cdot 10^{-8}\ m)}{62 \cdot (1.45 \cdot 10^{-10}\ m)}}$$

$$= 4.4 \cdot 10^{-18}\ mol\ s^{-1}$$

The rate constant of this diffusive current is

$$k = (4.4 \cdot 10^{-18}\ mol\ s^{-1}) \cdot (6.022 \cdot 10^{23}\ mol^{-1})$$

$$= 2.6 \cdot 10^{6}\ s^{-1}$$

In Hinzpeter's model of diffusion, the permeability of the pores to the diffusing species is the main factor affecting diffusion. Shell permeability is dependent on the shell surface area, the number of pores in the shell, pore length, and effective pore radius. The effective pore radius is dependent on the radius of the pore and the radius of the diffusing species. The shell permeability is

$$p \simeq \frac{N_P \pi (r_p - r_w)^2}{A} \frac{D}{\delta}$$

where $p$ is shell permeability in m $s^{-1}$, $N_P$ is the number of pores, $r_p$ is the pore radius in m, $r_w$ is the radius of the diffusing species in $m^2$, $A$ is the surface area of the shell in $m^2$, $D$ is the diffusion coefficient in $m^2$ $s^{-1}$, and $\delta$ is the pore length in m.

Hess *et al.*[6] phrases Hinzpeter *et al.*'s model in the context of a diffusive current dependent on the concentration of diffusing species and the diffusive conductance, which is in turn dependent on pore permeability. The rate constant of diffusion using Hinzpeter's model of diffusion is

$$k = NN_A \pi s^2 \frac{D}{\delta} C_0$$

where $k$ is the rate constant in $s^{-1}$, $N$ is number of pores, $N_A$ is Avogadro's number in $mol^{-1}$, $s$ is the effective pore radius in m, $\delta$ is pore length. The effective pore radius of the wild-type encapsulin in this scenario is equal to the difference in radius between the encapsulin pore (1.45 Å) and the ionic radius of $Tb^{3+}$ (1.2 Å). An average pore length of 11 Å is assumed, based on measurements from the crystal structure. All other factors shared between this calculation and the above calculation are kept constant. Based on this, the rate constant for $Tb^{3+}$ diffusion is

$$k = 62 \cdot (6.022 \cdot 10^{23}\ mol^{-1}) \cdot \pi \cdot (2.5 \cdot 10^{-11}\ m)^2 \cdot \frac{(6 \cdot 10^{-10}\ m^2 s^{-1})}{(1.1 \cdot 10^{-9}\ m)} \cdot (2.5 \cdot 10^{-1}\ mol\ m^{-3})$$

$$= 1.0 \cdot 10^4\ s^{-1}$$
